# Intrinsically disordered insert from SH2D2A rewires CD19 CAR signaling via Tyr290

**DOI:** 10.64898/2026.02.03.703491

**Authors:** Pawel Borowicz, Brian Christopher Gilmour, Hanna Chan, Ramakrishna Prabhu Gopalakrishnan, Timo Peters, René Platzer, Jacqueline Seigner, Johan Georg Visser, Hanna Kjelstrup, Aliki Popidou, Manal el Darwich, Ivan Abbedissen, Sebastian L. Andree, Stian Foss, Gustavo Antonio de Souza, Michael W. Traxlmayr, Vibeke Sundvold, Sébastien Wälchli, Johannes Huppa, Anne Spurkland

## Abstract

Chimeric antigen receptor (CAR) T cells have transformed cancer immunotherapy, yet their truncated or suboptimal intracellular signaling can limit therapeutic efficacy. To enhance proximal signaling of a CD19-targeted CAR, we systematically inserted short Lck-recruiting motifs derived from Lck-adaptor proteins into the CAR intracellular tail. Six candidate sequences from four adaptor molecules (SH2D2A, SKAP1, LAT, LIME), with a sequence from CD3ε, known to affect CAR functionality, as a positive control, were tested for expression and functional impact. Three CAR constructs (containing SH2D2A, LAT and LIME1 sequences respectively) displayed reduced surface expression, but only SH2D2A elicited a pronounced rewiring of CAR T cell phenotype following co-culture with CD19^+^ tumor lines. SH2D2A CAR T cells showed increased CD27 and CD56 expression and reduced expression of effector-associated mediators including granzyme B, IL-2, TNFα, and IFNγ. Through systematic mutagenesis and comparative phenotyping of SH2D2A CAR variants, we identified SH2D2A tyrosine 290 (Tyr290) as the critical residue mediating both the altered signaling phenotype and the low surface expression. Additionally, mutation of Tyr254 in the LIME1 CAR restored surface expression in Jurkat T cells, indicating insert- and context-dependent effects on receptor surface expression. Collectively, these results demonstrate that short, intrinsically disordered adaptor-derived sequences — and single tyrosine residues within them — can profoundly reprogram CAR signaling and expression.

## Introduction

Chimeric antigen receptors (CARs) uniquely combine the specificity of selected antibodies with the activation potential of T-cells. The first-generation CARs included an extracellular single-chain variable fragment (scFv) from an antibody, coupled with a transmembrane segment from CD4 and an intracytoplasmic signaling sequence derived from CD3ζ. However, these CAR T-cells demonstrated limited clinical effectiveness due to insufficient persistence in patients^1^.

The efficacy of CAR T-cells improved significantly with the introduction of co-stimulatory intracellular domains from receptors such as CD28 and 4-1BB, leading to the development of second-generation CARs. CARs with this enhanced intracytoplasmic sequence were approved by FDA for clinical use in treating B-cell lymphomas in 2017^2^ and multiple myeloma in 2021^3^, representing a major advancement in cancer immunotherapy.

Despite the remarkable advancements in CAR-T cell therapy, several early studies highlighted important aspects of CAR signaling. Antigen sensitivity may be significantly compromised by inefficient recruitment of intracellular signaling components^4^. Co-stimulatory molecule CD2 may be required for proper formation of the CAR-induced immunological synapse^5^. CARs may not require the adaptor protein LAT for synapse formation^6^.

Collectively, these findings suggested that truncated signaling pathways may indeed diminish the effectiveness of CAR-T cell physiological responses. The low proliferation and high exhaustion rates of the FDA-approved CAR-T cells, along with their inability to establish memory, can be linked to inadequate cellular signaling from the CAR^7^.

In response to this challenge, the initial successes of CAR T-cell therapy have spurred an influx of innovative CAR designs that often report improved functionality over standard CARs, particularly in addressing the T cell exhaustion seen with second-generation CARs.

Lck is a crucial cytosolic kinase that initiates TCR-triggered signaling in T cells. Lck is also the major kinase initiating phosphorylation of the CD3ζ tyrosine motifs of the CAR^8^. How to efficiently recruit Lck to the truncated signaling sequence of CAR is still a matter of intensive research^9^. Early experiments with chimeric receptors indicated that direct incorporation of Lck into CAR constructs significantly enhanced their signaling capacity^10^. Similarly, the addition of a single Lck interaction site in the CAR construct can amplify the antitumor responses of CAR T-cells^11^. Interestingly, although it may seem sufficient to simply couple Lck to the CAR construct^10^, some evidence suggests that removing Lck binding sites from CAR^12^ or Lck protein from CAR T cell^8^, can actually improve the anti-tumor properties of CARs. The mechanisms underlying this discrepancy remain poorly understood and require further exploration. In this study, we aimed to enhance the functionality of CARs by incorporating sequences from Lck adaptor proteins in the CAR intracytoplasmic tail.

Physiologically, subtle differences in recruitment mechanisms of molecules are mediated by a class of proteins known as adaptor proteins. These molecules do not possess intrinsic enzymatic activity; rather, they function as molecular scaffolds that facilitate signaling. Our recent review highlighted the role of adaptor proteins in modulating signaling cascades within immune cells^13^.

Based on this, we hypothesized that by modifying Lck recruitment through the strategic inclusion of Lck-binding motifs derived from Lck adaptor molecules, CAR T-cell function can be significantly affected. This may potentially allow for fine-tuning of CAR T-cell expression, antigen sensitivity, cytokine production, and cytotoxic responses.

Here we designed and tested CAR constructs that incorporated various Lck-recruiting motifs derived from LAT, LIME1, SKAP1, and SH2D2A^13^ to elucidate the requirements for effective Lck recruitment to CARs.

While evaluating the functionality of our novel CAR designs, we uncovered an Lck adaptor region which rewires CAR signaling as well as a previously unreported mechanism impacting CAR surface expression. Taken together, we show that exploring the functionality of novel CAR designs not only has the potential to improve immunotherapy but also may be a tool to identify the biological role of applied protein motifs.

## Results

### Lck adaptors’ sequences affect surface expression of CAR constructs

To address how variations in Lck recruitment to CARs affect CAR-T cell function, we delved into identifying Lck-adaptors known for their role in modulating Lck function. Through an extensive literature review, we identified 16 proteins that met our definition of an “adaptor molecule”^13^ implicated in interactions with Lck (Sup. Tab. 1). For compatibility with CAR design, these interactions needed to depend on Lck binding a short linear motif (SLIM) within the adaptor protein’s intrinsically disordered region (IDR) (Sup. Fig. 1). Of the 16 candidates, only four molecules contained a well-characterized Lck-binding SLIM and thus satisfied this criterion. In the cases of LAT and SH2D2A, two distinct motifs were identified, and we opted to incorporate both into our CAR constructs.

For inclusion in the CARs, the motifs were extended to include their conserved flanking sequences, in order to retain the complete interaction interface, while enabling future optimization should it enhance CAR function. Although CD3ε does not meet our criteria for an adaptor molecule due to its role as an anchored receptor component^13^, its RK motif recruits Lck via non-canonical binding by the Lck SH3 domain—a mechanism aligned with the objectives of this study. When included into CAR design, the RK-motif has been shown to enhance CAR functionality^11^. We therefore utilized the RK-motif derived from CD3ε as a benchmark control for Lck recruitment to CAR.

Our constructs were based on a second-generation, FDA approved CD19-CAR (CD28TM-CD28-CD3ζ)^14^ featuring a T2A self-cleaving peptide and a truncated CD34 (tCD34) as a marker for expression of the CAR^15^, assembled within a single lentiviral plasmid^16^. The sequences of selected Lck adaptors (SH2D2Aseq1, SH2D2Aseq2, SKAP1, LATseq1, LATseq2, LIME1, and CD3ε) (Fig. 1A) were inserted between CD28 and CD3ζ (Fig.1B).

**Figure 1.**
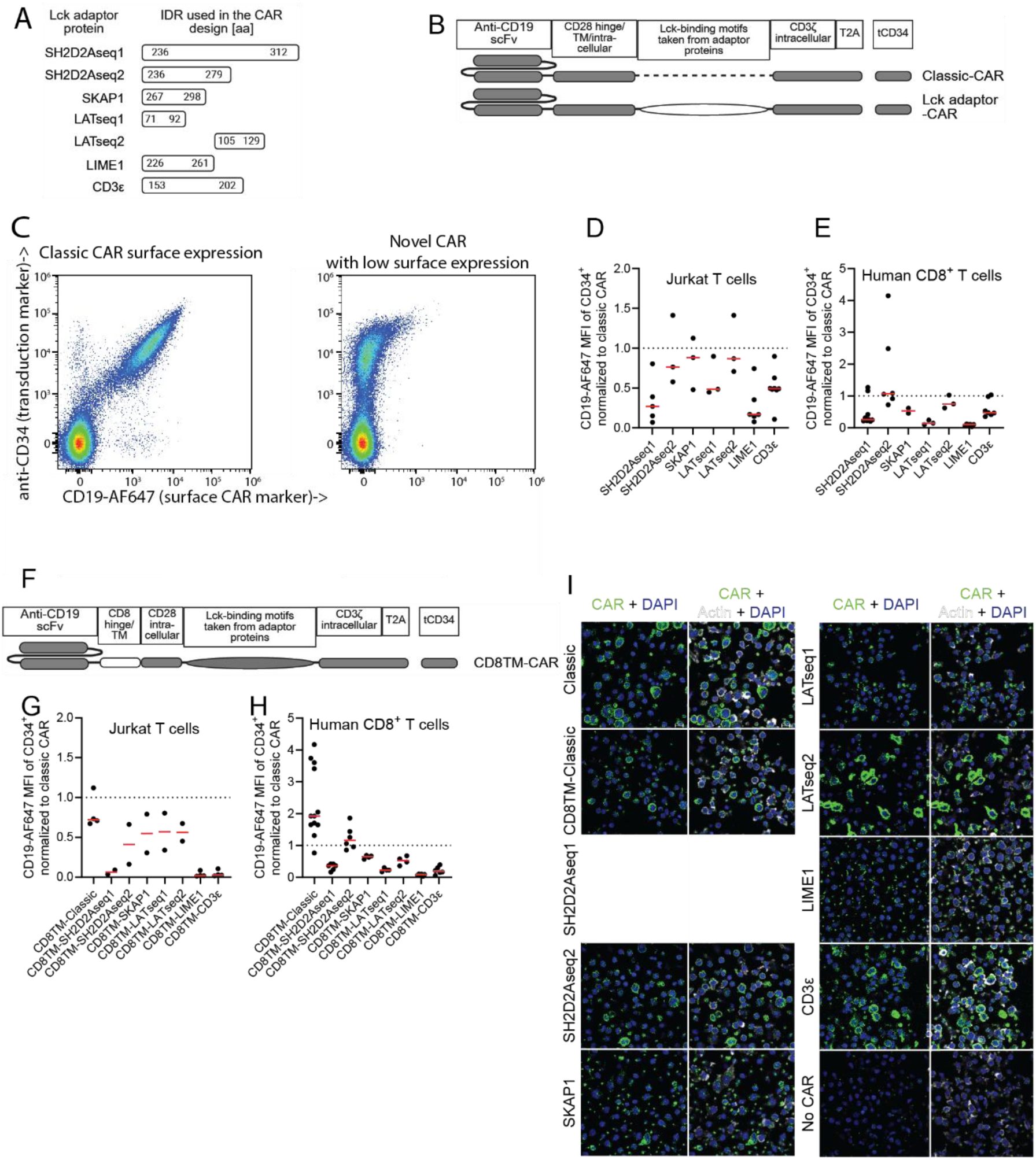
Lck adaptors’ sequences affect surface expression of CAR constructs. A) Schematic representation of Lck adaptors sequences used in this study. B) Schematic representation of CD19-CARs with inserted Lck adaptors sequences. C) Representative dot plots of two CD19-CAR constructs transduced into human CD8^+^ T cells which have either high or low surface expression as assessed with SuperFolder CD19 (surface expression) and anti-CD34 (transduction marker) staining. D) Summary graph of surface expression (MFI) of CD19-CARs with Lck adaptors sequences expressed in JE6.1 T cells. Each dot represents a separate experiment (n=3-8). All data was normalized to the classic CD19-CAR measured within the given experiment. E) As in D), MFI of indicated CARs expressed in human CD8^+^ T cells. Each dot represents a separate experiment (n=2-9) performed in n=2-5 donors. All data was normalized as in D). F) Schematic representation of CD19-CARs with inserted Lck adaptors sequences and CD28TM domain replaced with CD8TM. G) Summary graph of surface expression (MFI) of CD8TM CD19-CARs with Lck adaptors sequences expressed in JE6.1 T cells. Each dot represents separate experiment (n=2-4). All data was normalized to classic CD19-CAR measured within given experiment. H) As G), MFI of indicated CARs expressed in human CD8^+^ T cells. Each dot represents separate experiment (n=4-12) performed in n=1-5 donors. All data was normalized as in G). I) Immunofluorescent staining of CD34^+^ primary human CD8^+^ T cells transduced with CD19-CARs with Lck adaptors sequences. CARs were identified with anti-T2A antibody.

In order to validate our constructs, we transduced them into both Jurkat T cells (JE6.1) and primary CD8^+^ human T cells. After gating cells on tCD34 (Sup. Fig. 2A), we were able to assess the CAR surface expression with SuperFolder CD19^17^ (Fig. 1C). All constructs were similarly expressed in JE6.1 and primary CD8^+^ human T cells, as indicated by tCD34 expression (Sup. Fig. 2B-C). However, certain sequences incorporated into the CAR constructs significantly reduced the surface expression of the CARs in both JE6.1 (Fig. 1D and Sup. Fig. 2D) and primary CD8^+^ human T cells (Fig. 1E and Sup. Fig. 2E). In order to measure the total expression of CAR, we pulled down CAR protein from JE6.1 CD34^+^ CAR T cells lysates. Immunoblotting (Sup. Fig. 2F) revealed that reduced surface expression corresponds to reduced total protein expression.

Even CAR bearing the CD3ε sequence, which has progressed beyond the preclinical phase, displayed reduced surface expression compared to the unmodified second-generation CAR. This has also previously been noted^18–20^, while improved expression of a modified CD3ε-CAR construct was attributed to dimerization mediated by cysteines in the CD8 hinge region^20^. We thus exchanged the TM (transmembrane) and hinge domain of CD28 with those coming from CD8 in all our constructs (Fig. 1F). Surprisingly, this modification did not rescue the surface expression of the CARs in JE6.1 T cells (Fig. 1G and Sup. Fig. 2G) nor in CD8^+^ T cells (Fig. 1H and Sup. Fig. 2H). On the contrary, the replacement appeared to exacerbate the disparities in surface expression, despite unaltered expression of the constructs as indicated by tCD34 marker (Sup. Fig. 2I-J).

Imaging primary CD8^+^ human CD34^+^ CAR T cells using immunocytochemistry revealed that the intensity of intracellular CAR expression correlated with the corresponding surface expression (Fig.1I).

### Functional phenotype performance of human T cells with CARs with Lck adaptor sequences

Having confirmed that the novel CAR constructs were expressed in T cells, we assessed the functional significance of the inserted Lck-adaptor motifs in human primary CD8^+^ CAR T cells. CD8^+^ CAR T cells were cocultured for one week with four different cancer cell lines that either did or did not stably express tCD19-mCherry as the target (Fig. 2A). Samples were analyzed on days 2 and 7 using a comprehensive antibody panel and spectral flow cytometry, producing a semiomics dataset that tracked canonical T cell activation parameters over time and across target cell types. By comparing identical cancer cell lines with and without CD19 expression, we distinguished on-target from off-target effects. Because only ∼20% of T cells in each sample were CAR transduced—mimicking physiological heterogeneity—the untransduced T cells provided internal controls for within sample comparisons.

**Figure 2.**
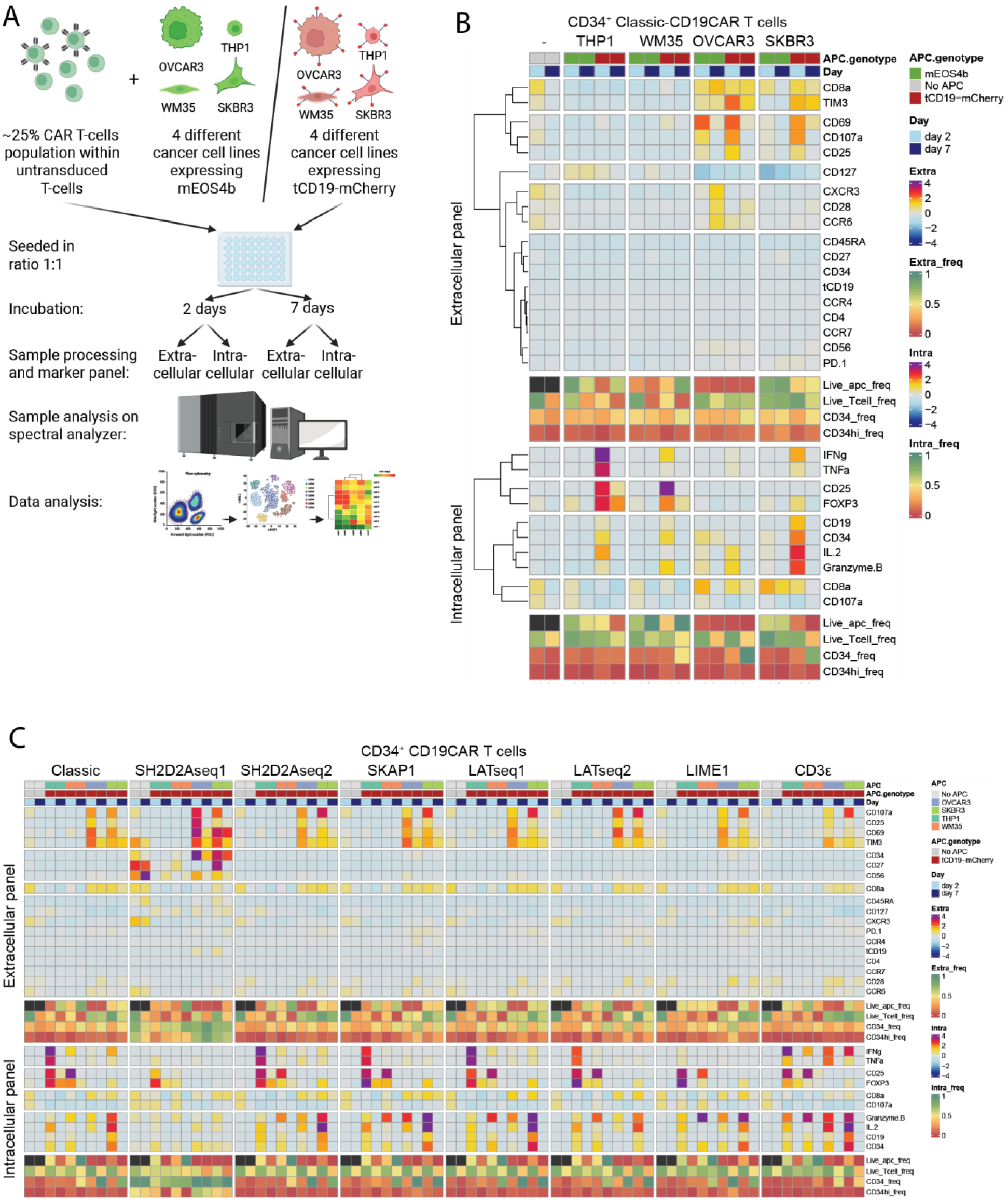
Functional phenotype performance of human T cells with CARs with Lck adaptor sequences. A) Schematic depiction of CAR T cells and target cells in a co-culture assay and its analysis with spectral flow cytometry. B) Heatmap presents mean normalized MFI values of various markers measured in CD19-CAR (classic) T cells from 2 donors incubated with the four different target cell lines expressing CD19 or not. C) Heatmap presents mean normalized MFI values of various markers measured in CD19-CAR (with Lck adaptor sequences) T cells from 2 donors incubated with the four different target cell lines expressing CD19.

To validate our CAR T cell activation assay, we first compared the response of Classic-CAR T cells exposed to CD19^+^ or CD19^−^ targets respectively. As expected, CD19^+^ cancer cell lines induced substantially greater activation of Classic-CAR T cells than their CD19^−^ counterparts (Fig. 2B). However, the magnitude and pattern of on-target activation was dependent on the target cell lines. Specifically: (1) THP1 induced IFNγ, TNFα, IL2, FOXP3, and intracellular CD25; (2) WM35 induced IFNγ, granzyme B, FOXP3, and intracellular CD25; (3) OVCAR3 increased surface CD69, CD25, CD107a, CD8, and TIM3, together with IL2 and granzyme B; (4) SKBR3 increased surface CD69, CD25, CD107a, CD8, and TIM3 and also induced IL2, granzyme B, IFNγ, and FOXP3. OVCAR3 cells elicited the most pronounced off-target activation (Fig. 2B). Untransduced (CAR^−^) T cells did not show target specific activation patterns and did not produce cytokines (Sup. Fig. 3A).

We then compared the response upon co-culture with CD19^+^ cancer cell lines of T cells expressing classic CAR with that of T cells expressing our novel Lck-adaptor CAR variants (Fig. 2C). With the exception of SH2D2Aseq1-CAR, the Lck-adaptor CAR constructs produced similar activation profiles that differed mainly in magnitude. Notably, relative to the other CARs the CD3ε-CAR elicited pronounced TNFα and IFNγ responses to OVCAR3 and SKBR3. SH2D2Aseq1-CAR T cells displayed a distinct phenotype characterized by elevated CD27 and CD56 expression, a markedly stronger response to OVCAR3 and SKBR3, and a markedly weaker response to THP1 and WM35. Cytokine and granzyme B induction/accumulation by SH2D2Aseq1 was comparatively low. Moreover, SH2D2Aseq1-T cells tended to upregulate CD27 and CD56 and showed nonspecific activation even when incubated with OVCAR3 or SKBR3 without CD19 (Suppl. Fig. 2B). Also at baseline, in the absence of target cells, SH2D2Aseq1-CAR T cells displayed an expression pattern distinct from the other CAR T cells including higher amounts of CD27, CD56, TIM3, CXCR3, and CCR6 (Fig. 2C).

### Certain phosphotyrosines in the CAR can affect its expression

While several of the CAR displayed significantly lower CAR surface expression (Fig. 1E), all the different CAR T cells were able to respond to target cells in an antigen specific manner (Fig. 2C and Sup. Fig. 3B).

To exclude the possibility that CAR surface expression might affect CAR T cell responsiveness, we investigated this issue further. We focused on CARs including the Lck adaptor SH2D2A^13^, since the length of the included sequence (SH2D2Aseq1 aa236-312 vs SH2D2Aseq2 aa236-279) seemed to correlate with a striking difference in the corresponding CAR surface expression (Fig. 1D-E and G-H). Given that the order and spacing of co-stimulatory domains in CARs can alter receptor function^21^, we first asked whether the placement of the Lck adaptor’s sequence within the SH2D2Aseq1-CAR structure influenced its expression (Fig.3A). Switching the order of the CD3ζ and adaptor sequence did not affect CAR surface expression nor overall expression of the constructs, irrespective of hinge region (CD28 or CD8), or cell type (CD8^+^ or JE6.1 T cells) (Fig.3B-C and Sup. Fig. 4A-D).

**Figure 3.**
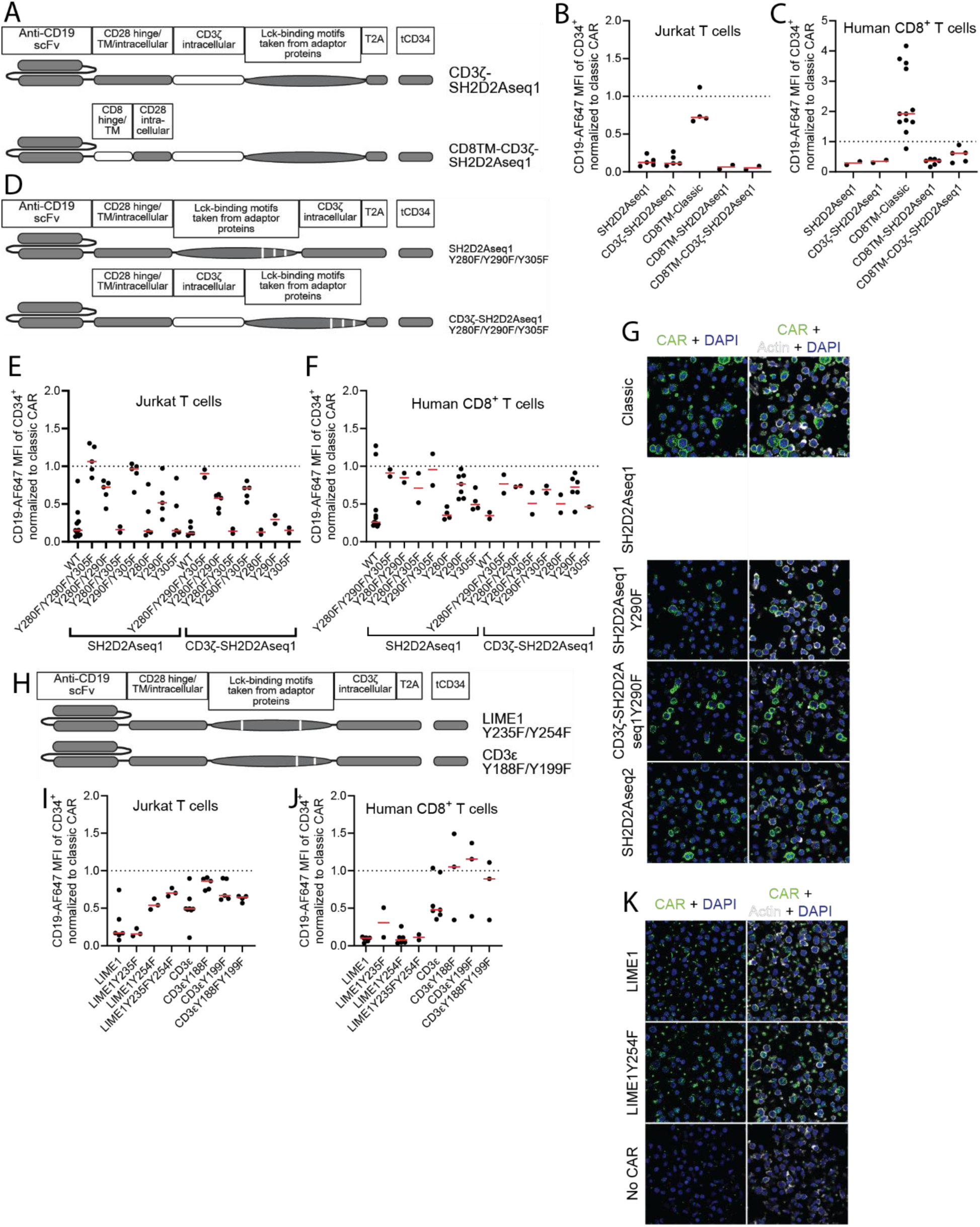
Certain phosphotyrosines in the CAR can affect its expression. A) Schematic representation of CD19-CARs with different transmembrane regions and with Lck adaptors sequences inserted after CD3ζ domain. B) Summary graph of surface expression (MFI) of CD19-CARs with Lck adaptors sequences after CD3ζ domain expressed in JE6.1 T cells. Each dot represents separate experiment (n=2-5). All data was normalized to classic CD19-CAR measured within given experiment. C) As in B), MFI of indicated CARs expressed in human CD8^+^ T cells. Each dot represents separate experiment (n=2-12) performed in n=2-5 donors. All data was as in B). D) Schematic representation of SH2D2Aseq1 CD19-CARs with mutated phosphotyrosines in all iterations including also CD3ζ domain switch. E) Summary graph of surface expression (MFI) of SH2D2Aseq1 CD19-CARs with mutated phosphotyrosines expressed in JE6.1 T cells. Each dot represents separate experiment (n=2-5). All data was normalized to classic CD19-CAR measured within given experiment. F) As in E), MFI of indicated CARs with mutated phosphotyrosines expressed in human CD8^+^ T cells. Each dot represents separate experiment (n=2-6) performed in n=2-4 donors. All data was normalized as in E). G) Immunofluorescent staining of CD34^+^ primary human CD8^+^ T cells transduced with CD19-CARs with Lck adaptors sequences. CARs were identified with anti-T2A antibody. H) Schematic representation of LIME1 and CD3ε CD19-CARs with mutated phosphotyrosines in all iterations. I) Summary graph of surface expression (MFI) of LIME1 and CD3ε CD19-CARs with mutated phosphotyrosines expressed in JE6.1 T cells. Each dot represents separate experiment (n=3-5). All data was normalized to classic CD19-CAR measured within given experiment. J) As in I), indicated CARs expressed in human CD8^+^ T cells. Each dot represents separate experiment (n=2-7) performed in n=2-5 donors. All data was normalized as in I). K) Immunofluorescent staining of CD34^+^ primary human CD8^+^ T cells transduced with CD19-CARs with Lck adaptors sequences. CARs were identified with anti-T2A antibody.

The SH2D2Aseq2-CAR construct that expressed well on the cell surface contained only the N-terminal proline-rich region, while the SH2D2Aseq1-CAR with less pronounced cell surface expression also contained three potential phosphotyrosines (pTyr) critical to SH2D2A’s function. To parse the effects of these residues on surface expression, we mutated each of the three tyrosines to phenylalanine (Phe, mimicking a non-phosphorylated residue) in all possible combinations (Fig. 3D). Only CAR constructs featuring the Y290F mutation displayed restored surface expression in JE6.1 T cells irrespective of localization in the CAR-construct (Fig. 3E and Sup. Fig. 4E-F). In CD8^+^ T cells, this effect was less striking, but the pattern was nearly identical (Fig. 3F and Sup. Fig. 4G-H). Mutating tyrosines did not change the constructs’ overall expression as indicated by tCD34 marker (Sup. Fig. 4I-J). In order to measure the total expression of CAR, we pulled down CAR protein from JE6.1 CD34^+^ CAR T cells lysates. Immunoblotting (Sup. Fig. 4K) revealed that restoration of surface expression corresponds to restoration of total protein expression.

Two of the three additional CARs with reduced surface expression, LIME1 and CD3ε, contained potential phosphotyrosine sites as well (LATseq1 does not contain any tyrosines). To explore whether also some of these tyrosines affected surface expression of the CAR, we performed tyrosine-to-phenylalanine mutations in the LIME1 and CD3ε constructs (Fig. 3H). Surface expression of LIME1Y254F-CAR was fully restored in JE6.1 T cells (Fig.3I and Sup. Fig. 4L) but not in primary CD8^+^ T cells (Fig.3J and Sup. Fig. 4M). Tyrosine mutations did not improve CD3ε-CAR surface expression in neither cells (Fig.3I-J and Sup. Fig. 4L-M). Mutating tyrosines in the LIME1 and CD3ε constructs did not change the constructs expression as indicated by tCD34 marker (Sup. Fig. 4N-O). Thus, elimination of a potential phosphotyrosine in novel designs of CD19-CAR could sometimes but not always be a means to restore their surface expression.

### Mutation of Tyr290 in SH2D2Aseq1-CAR abolishes its ability to rewire CAR signaling

Since we found that some tyrosine residues control surface expression of certain CAR designs, we proceeded to test whether these tyrosines also affected CAR T cell function. When our Lck adaptors CAR T cells were co-cultured with OVCAR3-tCD19-mCherry target cells, they formed synapses (Fig. 4A) and the occurrence of synapses mostly corresponded to surface expression of the given CAR (Fig. 1D). Therefore, mutating Tyr290 in SH2D2Aseq1-CARs and Tyr254 in LIME1-CAR correlated with increased occurrence of synapses (Fig. 4B). This suggested that the mutated CAR variants could display an enhanced function in co-culture assay.

**Figure 4.**
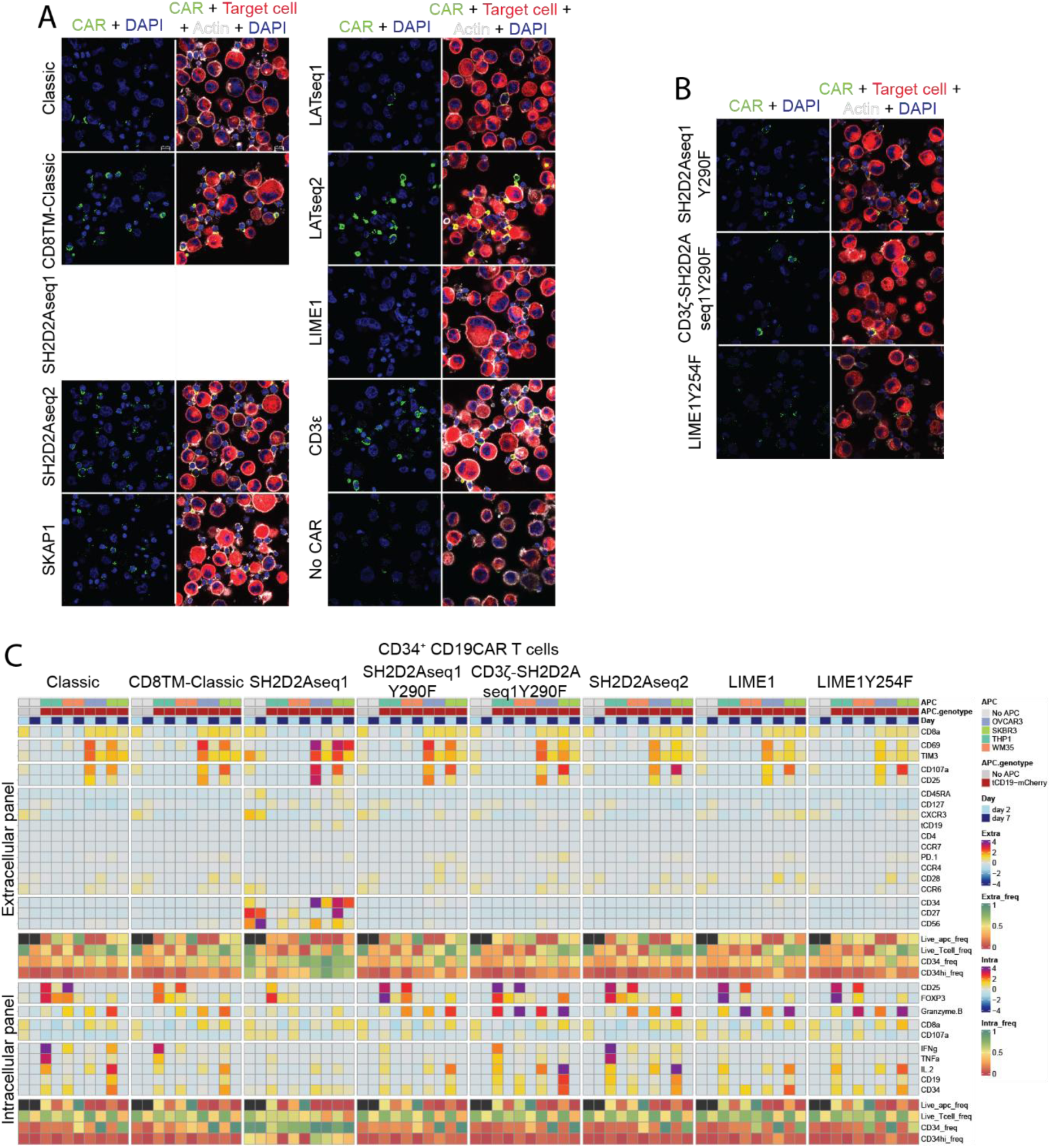
Mutation of SH2D2A Tyr290 in CD19-CAR with SH2D2Aseq1 abolishes its rewired signaling. A) Immunofluorescent staining of CD34^+^ primary human CD8^+^ T cells transduced with CD19-CARs with Lck adaptors sequences and incubated with OVCAR3-tCD19-mCherry target cell line. CARs were identified with anti-T2A antibody. B) Immunofluorescent staining of CD34^+^ primary human CD8^+^ T cells transduced with CD19-CARs with SH2D2A and LIME1 sequences and incubated with OVCAR3-tCD19-mCherry target cell line. CARs were identified with anti-T2A antibody. C) Altered CAR T cells with SH2D2A or LIME1 sequences vs target cell co-culture assay. Heatmap presents the comparison between various altered CD19-CAR (with SH2D2A or LIME1) T cells incubated with the four different target cell lines expressing CD19.

Thus, we selected three CAR variants for functional testing in the same coculture assay as used in Fig. 2: SH2D2Aseq1Y290F, CD3ζSH2D2Aseq1Y290F, and LIME1Y254F. Each construct was cocultured for 2 or 7 days with the four cancer cell lines, with or without CD19 expression, and CAR T cell activation phenotypes were assessed by spectral flow cytometry (Fig. 4C).

Surprisingly, all the tested tyrosine to phenylalanine mutated CARs produced T cell activation phenotypes similar to the classic CAR (Fig. 4A). The most notable change in marker expression was observed in SH2D2Aseq1Y290F-CAR T cells, relative to the unmutated SH2D2Aseq1-CAR T cells. When these tyrosine to phenylalanine mutated CAR T cells were cocultured with CD19^−^ cancer cells, their marker profiles also reverted to resemble the classic CAR (Sup. Fig. 5A).

### SH2D2A pTyr290 is not recognized by phosphotyrosine binding proteins

The three SH2D2A tyrosines (280, 290, and 305) are highly conserved^13^ (Sup. Fig. 6A-B), and we have previously found that Lck both phosphorylates and interacts via its SH2 domain with all three tyrosines^22^. The striking effect of Tyr290 in SH2D2Aseq1 on CAR T cells activation led us to revisit the specific role of this tyrosine in regulating SH2D2A function.

When intact or mutated SH2D2A cDNA was exogenously over-expressed in anti-CD3 activated Jurkat T cells, Tyr290 was found to be necessary for its maximal tyrosine phosphorylation (Fig. 5A-B). To further explore the effect of phosphorylation of Tyr290 we generated endogenously expressed SH2D2A Y290F and Y290E (Glu, mimicking a phosphorylated residue) knock-in Jurkat T cells using CRISPR/Cas9 genome editing^23^. Mutants were pre-activated with PMA/IO for 16 hours to induce SH2D2A expression, and stimulated with anti-CD3, treated with PV (pervanadate) or left untreated, followed by Western blot analysis of the presence of SH2D2A. In lysates from untreated cells, SH2D2A Y290F migrated similarly to wild-type (WT), while SH2D2A Y290E migrated similarly to PV-treated WT SH2D2A (Fig. 5C). Furthermore, the migration patterns of both mutants were unaffected by PV treatment, supporting the idea that the higher molecular weight band of SH2D2A represents phosphorylation on Tyr290^22^.

**Figure 5.**
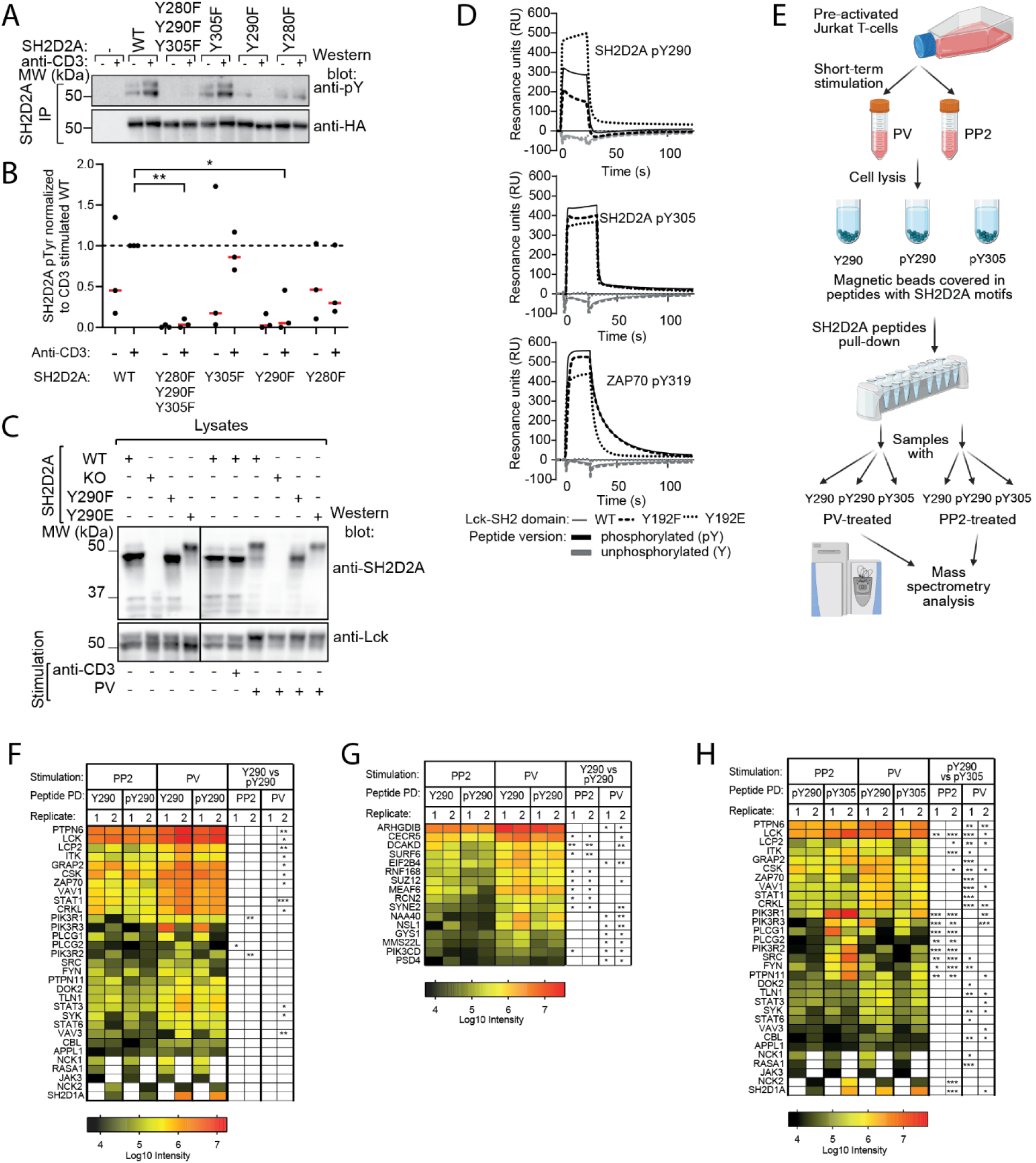
SH2D2A Tyr290 is not recognized by phosphotyrosine binding proteins. A) Jurkat cells transiently transfected with the indicated plasmid constructs were stimulated with anti-CD3 antibody for 2.5 min. SH2D2A was immunoprecipitated and detected by immunoblotting with anti-phosphotyrosine (pTyr) and anti-HA antibodies. One representative immunoblot is shown (n = 3). B) Summary graph of A). Each point represents an independent experiment (n = 3). Data were normalized to the anti-CD3–stimulated SH2D2A wild-type (WT) sample within each experiment. C) SH2D2A WT and SH2D2A knock-in mutated JTag cell lines pre-activated with PMA/IO, were treated with PV (5min), with anti-CD3 antibody (2min) or left untreated before being lysed. Samples were immunoblotted with the indicated antibodies. One representative immunoblot is shown (n=3). D) Peptides representing SH2D2A pTyr290, pTyr305, ZAP70 pTyr319 and their unphosphorylated versions were probed with purified Lck-SH2 domains representing Lck WT, Y192F and Y192E mutated versions using SPR. Each graph shows the binding profiles of one of the Tyr motifs, both phosphorylated and unphosphorylated. The Lck-SH2 domains’ concentrations were as follows: 5μM for SH2D2A Tyr290 peptides, 4μM for SH2D2A pTyr305, 5μM for SH2D2A Tyr305, 2,5μM for ZAP70 peptides. E) Schematic depiction of SH2D2A peptide pull down performed in JTag T cells and analyzed with mass spectrometry (n=2). F) Heatmap presents median intensity values from three technical replicates (in two biological replicates) of MS analysis of proteins interacting with SH2D2A Tyr290 and pTyr290 peptides respectively, filtered for proteins containing SH2 or PTB domains. Significant differences between pairwise comparisons within each experiment are indicated with stars as explained below. G) Heatmap presents MS results (as in B) for proteins showing significant enrichment in either SH2D2A Tyr290 or pTyr290 PD samples which was consecutive for both replicate experiments in given condition. H) Heatmap presents MS results (as in B) from SH2D2A pTyr290 and pTyr305 peptide PD, filtered for proteins containing SH2 or PTB domains. The color intensity bar is shown below each corresponding heatmap. Statistically significant comparisons are shown on the heatmaps accordingly: * - p-value <0.05; ** - p-value <0.01; *** - p-value <0.001.

Our previous work suggested that Tyr290 control over SH2D2A phosphorylation might come from preferred binding to the kinase, since phosphorylation of Lck-SH2 on Tyr192 alters its binding preference from SH2D2A pTyr305 to pTyr290^24^. To confirm this, we applied surface plasmon resonance (SPR) to assess the binding of wild-type (WT), Y192E, or Y192F Lck-SH2 domains to SH2D2A peptides (Fig. 5D). ZAP70 pTyr319, a known target of Lck-SH2^25^, was used as a control. Our SPR data revealed that the WT and Y192F Lck-SH2 domains bound both ZAP70 pTyr319 and SH2D2A pTyr305 peptides with similar affinities, reinforcing pTyr305’s role as a canonical SH2-binding motif. In contrast, and in accordance with previous analysis by calorimetry^24^ the Y192E Lck-SH2 domain, which mimics Lck phosphorylation on Tyr192, displayed a stronger preference for binding SH2D2A pTyr290 (Fig.5D).

To further dissect the role of SH2D2A Tyr290 as a binding motif, we established and compared Tyr290 and Tyr305 interactomes in Jurkat T cells. Briefly, biotinylated peptides corresponding to SH2D2A Tyr290, pTyr290, and pTyr305 were used to pull-down (PD) interacting proteins in lysates from Jurkat T cells (Fig. 5E) followed by mass spectrometry (MS) to identify SH2D2A associated proteins. Since SH2D2A is upregulated in activated T cells^26^, JTAg T cells were pre-activated with PMA/ionomycin (PMA/IO) for 16 hours before the experiments. Additionally, recognizing that some protein-protein interactions (including Lck binding to SH2D2A pTyr290) are phosphorylation-dependent^24^, prior to lysis, cells were pre-treated with either pervanadate (PV) to maximize protein phosphorylation or the Src inhibitor PP2 to minimize it.

We first assessed the presence of SH2 or PTB domain-containing proteins in the MS dataset and found no significant differences in the ability of SH2D2A Tyr290 vs. pTyr290 peptides to pull down members of this group of proteins (Fig. 5F). To extend our analysis, we searched the MS dataset for proteins whose interaction with Tyr290 peptides was significantly enriched under similar experimental conditions in both replicates (Fig. 5G). Sixteen proteins were identified that associated more frequently with Tyr290 than with pTyr290 peptides. Notably, no proteins exhibited preferential binding to pTyr290 peptides.

Next, we compared the binding profiles of pTyr290 and pTyr305 peptides to SH2 or PTB domain-containing proteins (Fig. 5H). In PP2-treated samples, binding of phosphatidylinositol-3-kinase subunits (PIK3R1-3), phospholipase C gamma (PLCG1, PLCG2), Src family kinases (LCK, FYN, SRC), and protein tyrosine phosphatase (PTPN11) was significantly enriched with pTyr305 peptides. However, in PV-treated cells, several proteins, including protein tyrosine phosphatase (PTPN6), tyrosine kinases (LCK, CSK, SYK), adaptor proteins (CRKL, LCP2), cytoskeletal protein (TLN1), guanine nucleotide exchange factor (VAV1), and E3 ubiquitin-protein ligase (CBL), showed increased association with pTyr290 peptides. Despite this, the binding of these proteins to Tyr290 and pTyr290 peptides was generally similar (Fig. 5D). Some of the PD hits were validated using western blotting in both Jurkat T cells (Sup. Fig. 6C) and CD4^+^ primary human T cells (Sup. Fig. 6D). To exclude the possibility of co-precipitation via LCK interaction, we further confirmed these results in LCK knockdown Jurkat T cells (Sup. Fig. 6E).

### Mutation of Tyr290 in SH2D2Aseq1-CAR restores its colocalization with Lck

Our results so far indicate that Tyr290 serves as a unique motif directly linked to SH2D2A function. To further explore the role of this particular Lck adaptor sequence for CAR function, we evaluated Lck recruitment to all our novel CAR designs by immunocytochemistry of CAR-expressing JE6.1 T cells (Fig. 6A). All CARs localized to the plasma membrane; notably, the CAR–SKAP1 construct also accumulated in an intracellular compartment, most likely the Golgi apparatus. Although the Classic-CAR showed strong co-localization with Lck, this pattern was not universally observed. In JE6.1 CD34+ CAR T cells, both the SH2D2Aseq1 and SKAP1 constructs failed to co-localize/co-express with Lck. In these cells CAR signal intensity was highest where Lck signal was lowest (Fig. 6A).

**Figure 6.**
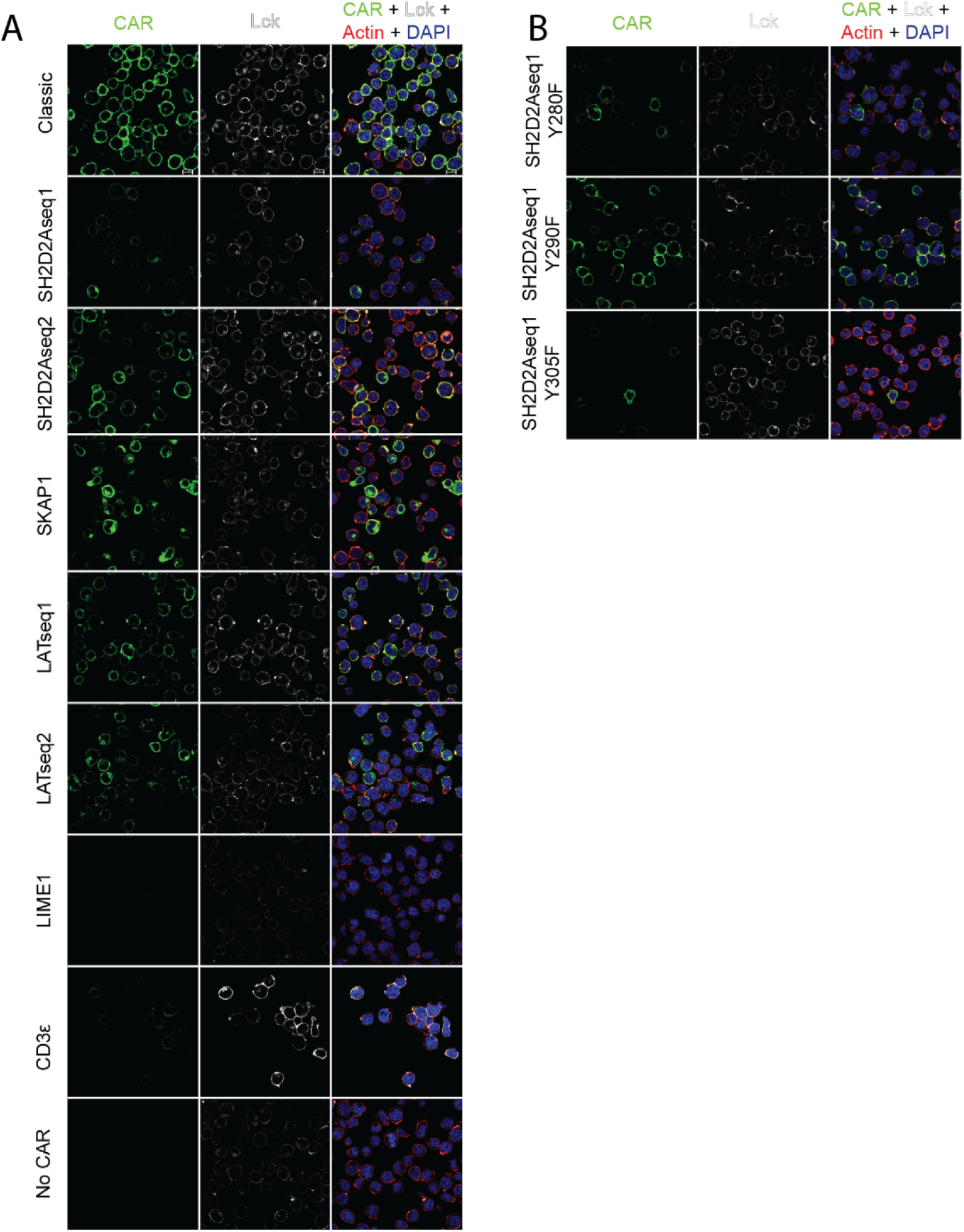
Lck expression (colocalization) in CARs with Lck adaptor sequences varies. A) Immunofluorescent staining of CD34^+^ JE6.1 T cells transduced with CD19-CARs with Lck adaptors sequences. CARs were identified with anti-T2A antibody. B) Immunofluorescent staining of CD34^+^ JE6.1 T cells transduced with CD19-CARs with mutated SH2D2A sequences. CARs were identified with anti-T2A antibody.

To probe the interaction further, we imaged T cells expressing tyrosine mutated SH2D2Aseq1-CAR. We found that the Y290F mutation restored the CAR co-localization with Lck (Fig. 6B). These results support the hypothesis that Tyr290 regulates SH2D2A function via Lck and help explain the pronounced functional rewiring observed for the SH2D2Aseq1-CAR.

## Discussion

While exploring the effect of adding Lck adaptor sequences to CAR constructs, we found that CAR surface expression may be impacted by specific tyrosine residues. One highly conserved tyrosine in SH2D2A, also rewired the activation of the CAR T cells, pointing to the functional relevance of this tyrosine not only in the CAR but also in the intact Lck adaptor. Our co-culture assay, combined with longitudinal phenotypic readouts, provided us with a cost-effective, straightforward, and ethically attractive entry point to preclinical studies of novel CAR’s design function.

The CAR’s design can control CAR surface expression in two ways: through transcription (transgene promoters and RNA structure) or through protein stability (receptor internalization, degradation, and recycling)^27^. Although acknowledged as an important mechanism, the latter is much less understood. Therefore, determining how single tyrosines are involved in these processes might not be trivial.

Many improvements to CAR-T cell design focus on enhancing the intracellular components, to best mimic the physiological response of T-cells activated through T-cell receptor (TCR). However, the rationale for selecting specific sequences to bolster signaling is frequently inadequate and relies on educated guesses. This may lead to a practice in which novel designs incorporating chosen protein sequences are dismissed if they do not manifest on the cell surface, as demonstrated in recent studies^28^. Such practice risks overlooking numerous potential CAR designs that significantly enhance CAR T-cell functionality but were discarded due to low surface expression.

CAR surface expression remains one of the most neglected yet critical factors influencing the success of CAR therapy^27^. To advance efforts to improve CAR designs, it is thus essential to understand the molecular features that determine the surface expression of CARs, particularly concerning the composition of endodomains. The role of endodomains in controlling CAR surface expression has been extensively studied in the context of CARs containing the CD3ε domain, though with many contradictory findings. For example, Velasco Cárdenas et al.^20^ demonstrated that the surface expression of CD3ε CARs depends strongly on dimerization, mediated by cysteines in the CAR’s hinge region derived from CD8. On the other hand, their attempt to restore surface expression of CD3ε CARs by removal of the endoplasmic reticulum (ER) retention motif (NQRRI) was unsuccessful. Xu et al.^18^ attributed the reduced surface levels of CD3ε CARs to faster CAR endocytosis. To address this, they removed a 27-amino acid interval between the basic residue-rich stretch and the ITAM, particularly targeting the proline-rich motif known to regulate TCR endocytosis and degradation. However, replacing this interval with six glycines only marginally improved surface expression. Wang et al.^19^ also suggested that CD3ε CAR surface expression was reduced due to ER self-retention. They successfully restored expression by replacing the CD28 signaling domain with 4-1BB or by removing specific fragments, such as GRQRGQNKER, YEPIRKGQRDLYSGL, or NQRRI, which shared a common feature: a significantly higher proportion of basic amino acids compared to acidic ones relative to other CD3 chains. Notably, our CD3ε CAR construct already lacked the NQRRI motif at the C-terminus, distinguishing it from those described in these studies.

For two of our Lck-adaptor CAR constructs (SH2D2Aseq1 and LIME1) we found that mutating a single highly conserved tyrosine to phenylalanine rescued surface expression of the CAR. To explore the significance of these tyrosines for CAR surface expression, we considered several possibilities. SH2D2A Tyr290 shares similarities with the C-terminal EFYA motif of syndecans that binds to PDZ domain-containing proteins^29^. Our mass spectrometry data revealed weaker binding to pTyr290 compared to unphosphorylated Tyr290. While this might support a mechanism involving PDZ domain binding, the Tyr290 motif is not located at the C-terminus (as is typical for PDZ-recognized motifs).

Tyrosines are also known to serve as sorting signals that direct transmembrane proteins to specific cellular compartments^30^. While the SH2D2A Tyr290 (QPIAFYAMG) and LIME1 Tyr254 (PLENVYESI) sequences do not fully match known sorting signal motifs, they bear some resemblance to the YF/Y motif utilized by HIV-1/SIV Nef for the endocytosis and degradation of CD4 and MHC-I^31^.

One other hypothesis for how tyrosines affect CAR surface expression, is that tyrosines may act as molecular switches that upon phosphorylation alter the conformation of the polypeptide in which they reside^13^. This could expose other non-tyrosine motif in the sequence, which control surface expression.

While the challenge of novel CARs lacking surface expression is recognized, to our knowledge, we are the first to demonstrate that single AA substitutions in the novel endodomain can resolve this issue. To assess the generality of this observation, it is however crucial to validate this phenomenon in greater detail and explore its application across other CARs with reduced surface expression. Especially in the light of data showing that CAR surface expression level might not be defining its usability. Wang et al. showed that less expressed CARs outperform higher expressed CAR designs by reducing cytokine release and thus improving safety and persistence of therapy^19^. Here, expanding on this idea, we show that even CAR T cells with undetectable surface expression of the CAR become activated in an antigen specific and functional manner. These data indicate that the prevailing paradigm of achieving high CAR expression for successful CAR therapy may be misguided or incorrect.

While alterations in CAR designs are primarily performed with the aim of possible clinical utility, our findings also demonstrate that their value in advancing basic research should not be overlooked. Incorporating sequences from signaling molecules with unknown functions into CARs offers an effective strategy for uncovering phenotypic effects and roles of these sequences or proteins. Using this approach, we uncovered novel insights into the T cell adaptor protein SH2D2A. Specifically, Tyr290 within the protein’s IDR emerged as a critical residue that determines the rewired functional phenotype of SH2D2Aseq1-CAR. Taken together with existing evidence implicating Tyr290 in SH2D2A regulation, our data support that this residue might be a principal mediator of SH2D2A function. That said, the physiological role of SH2D2A in T cells remains incompletely resolved, and further mechanistic and in vivo studies will be required to define its function definitively.

Although we identified a novel CAR design that produces a markedly rewired T cell phenotype, fully characterizing its importance for CAR T cell function will require additional experiments and may hinge on still uncharacterized functions of SH2D2A. Of particular interest is the unexpected combination of reduced cytokines and granzyme B expression together with enhanced target-cell killing. One hypothesis is that these CAR T cells employ alternative cytotoxic pathways, such as FASL-mediated killing, which requires further investigation. Other open questions include the mechanism underlying their (potentially trogocytic^32^) massive increase in CD34 expression and CD34^+^ population, and the basis for their divergent responses to different cancer cell lines. The latter behavior could reflect the contribution of mechanotransduction for this CAR design: THP1 and WM35 are weakly or not adherent, whereas OVCAR3 and SKBR3 are strongly adherent, and adhesion-dependent forces can profoundly influence T cell activation. Another interesting lead is the indication that the SH2D2Aseq1-CAR may signal independently of Lck. Prior work has shown that routing CAR signaling through Fyn rather than Lck can improve therapeutic performance^8^. Thus, altered kinase usage may contribute to the rewired phenotype observed here.

Our results revealed a clear discrepancy in CAR surface expression between JE6.1 T cells and primary CD8^+^ T cells. As JE6.1 is a widely used cell line for investigating novel CAR designs, this discrepancy may present significant challenges when transitioning to primary human T cells. The most striking difference was observed with CARs encoding the LIME1-Y254F mutation, which restored surface expression exclusively in JE6.1 T cells. Although we cannot entirely rule out the possibility that this effect is influenced by the CD4^+^ T-cell-like phenotype of JE6.1 cells (originally derived from CD4^+^ T cells), the most plausible explanation lies in the mutations that cause signaling abnormalities in JE6.1 cells—most notably, the constitutive activation of PI3K^33^.

Overall, our findings suggest that specific tyrosine residues can significantly influence CAR surface expression when incorporated into the construct’s endodomain. However, the precise mechanisms and rules governing how this amino acid residue, and potentially others, affect this process remain unclear. To successfully integrate novel signaling sequences into CARs, it is crucial to understand the “sequence grammar” that dictates surface expression. Therefore, further investigation into the role of tyrosines and the identification of other amino acids involved in this phenomenon is essential. Additionally, demonstrating that aiming for high level of CAR surface expression might not be relevant for its therapeutical goal is equally significant. More in-depth investigation of these aspects may offer valuable insights into CAR physiology and pave the way for innovative CAR designs. Ultimately, our novel CAR design (SH2D2Aseq1-CAR) with rewired functional capabilities should be suitable for consideration in pre-clinical trials.

## Methods

### Antibodies and reagents

Primary antibodies used were: anti-Lck (clone 3A5, Santa Cruz Biotechnology, RRID:AB_627880; clone IF6, a gift from Joseph B. Bolen), anti-Plcγ (1249, Santa Cruz Biotechnology, RRID:AB_632202), anti-CD3ζ (clone 6B10.2, Santa Cruz Biotechnology, RRID:AB_627020), anti-phosphotyrosine (clone 4G10, Upstate Biotechnology, RRID:AB_309678), anti-GAPDH (clone 6C5, Chemicon, RRID:AB_2107445), anti-PI3-kinase, p85 (clone 04-403, EMD Millipore, RRID:AB_673109), anti-SHP1 (clone C-19, Santa Cruz Biotechnology, RRID:AB_2173829), anti-SHP2 (C-18, Santa Cruz Biotechnology, RRID:AB_632401), anti-Itk (clone 2F12, Santa Cruz Biotechnology, RRID:AB_627834), anti-CD3ζ pY142 (clone K25-407.69, BD Biosciences, RRID:AB_647307), anti-CD3ε (clone OKT-3, American Type Culture Collection, RRID:AB_2073169), anti-T cell receptor (clone C305, a gift from Arthur Weiss, RRID:AB_417390), anti-CD28 (clone CD28.2, BD Biosciences, RRID: AB_396068), anti-SH2D2A antibodies raised against synthetic peptides of SH2D2A [1, 40] or the recombinant C terminus of SH2D2A (a gift from Virginia Shapiro). The secondary reagents and conjugated antibodies used were: HRP conjugated light chain specific mouse anti-rabbit (RRID:AB_2339146), HRP conjugated goat anti-mouse (H+L) (RRID:AB_10015289) and HRP conjugated goat anti-rabbit antibodies (RRID:AB_2307391) (Jackson ImmunoResearch). For stimulation or inhibition of cells: phorbol 12-myristate 13-acetate (PMA), ionomycin (IO), sodium orthovanadate, hydrogen peroxide, Src inhibitor PP2 (all Sigma-Aldrich). Peptides used were: biotin ZAP70 Tyr^319^ (Biotin-VYESPTyrSDPEE-NH2), biotin ZAP70 pTyr^319^ (Biotin-VYESPpTyrSDPEE-NH2)^25^ were from ProteoGenix (Schiltigheim, France), biotin SH2D2A Tyr^290^ (Biotin-QPIAFTyrAMG-NH2), biotin SH2D2A pTyr^290^ (Biotin-QPIAFpTyrAMG-NH2), biotin SH2D2A Tyr^305^ (Biotin-APSNITyrVEVED-NH2), biotin SH2D2A pTyr^305^ (Biotin-APSNIpTyrVEVED-NH2)^24^ were from Gencust (Dudelange, Luxembourg). All peptides were HPLC purified to at least 95% purity.

### Cell cultures

Human peripheral CD4^+^ and CD8^+^ T cells were isolated by positive selection from PBMC of anonymous healthy donors from the Norwegian Blood Bank (regional ethical committee project number 211844) by Dynabeads® CD4 Positive or CD8 Positive Isolation Kits (both ThermoFisher) as per the manufacturer’s protocol. Isolated cells were then used for downstream assays and cultured in AIM V media (with L-glutamine, 50μg/mL streptomycin sulfate, and 10μg/mL gentamicin sulfate) supplemented with 10% fetal calf serum (FCS), 1 mM sodium pyruvate, 1 mM non-essential amino acids, 1 mM HEPES buffer, 2 mM GlutaMAX (all from GIBCOBRL®, ThermoFisher Scientific), 50µM β-mercaptoethanol (Sigma-Aldrich), and 50-200 U/mL human recombinant IL-2 (ThermoFisher).

JE6.1^34^, JTAg (Jurkat T cells)^35^, THP1 (ATCC), OVCAR3, SK-BR-3, WM35 (a kind gift from prof. Erik Dissen, cell lines were authenticated at the Oslo University Hospital cell typing core facility^36^), and all CRISPR/Cas9 mutated or stably transduced cell lines were cultured in RPMI-1640 (with glutamine) supplemented with 10% fetal calf serum (FCS), 1 mM sodium pyruvate, 1 mM non-essential amino acids, 1 mM HEPES buffer, 2 mM GlutaMAX, 100 units/mL penicillin and 100 µg/mL streptomycin (all from GIBCOBRL®, ThermoFisher Scientific) and 50µM β-mercaptoethanol (Sigma-Aldrich).

Lentix (HEK293T cells, a kind gift from Dr. Coen Campsteijn) were cultured in DMEM (with high glucose, sodium pyruvate, and GlutaMax) supplemented with 10% fetal calf serum (FCS), 100 units/mL penicillin, and 100 µg/mL streptomycin (all from GIBCOBRL®, ThermoFisher Scientific).

All cells were cultured at 37°C in a humidified atmosphere at 5% CO2.

### Transient transfection

Jurkat cells were transfected by electroporation with a BTX electroporator (Genetronix). 10−15×10^6^ cells and 2–5 µg DNA were resuspended in 400 µl RPMI-1640 medium without antibiotics supplemented with 5% FCS and transfected at 240V for 25 ms. Cells were harvested 18–24 hours post-transfection.

### Plasmids for lentiviral transduction

All our plasmids dedicated for lentiviral transduction were assembled in pENTR vector (a kind gift from Dr. Sébastien Wälchli and Dr. Emmanuelle Benard) and transferred into pLX (Addgene #113668) vector via Gateway cloning system (Invitrogen, Thermofisher). Constructs were assembled with restriction enzyme cloning, Gibson Assembly^37^ (New England Biolabs), and/or megaprimer PCR^38^. Multiple point mutations were introduced into the pENTR-CD19-CAR-SH2D2A plasmid by QuickChange mutagenesis using Pfu-Turbo DNA polymerase (Agilent).

To generate our basic CAR model we merged CD19-CAR (FMC63 clone) (CD28TM-CD28-CD3ζ) sequence^14^ (a kind gift from Dr. Sébastien Wälchli and Dr. Emmanuelle Benard), via a viral T2A cleavage sequence, with a truncated CD34 receptor as a transduction marker^15,16^ (a kind gift from Dr. Sébastien Wälchli). Lck adaptor proteins (SKAP1, LATseq1, LATseq2, and LIME1) and CD3ε were cloned from a cDNA library (peripheral human CD8^+^ T-cells), while SH2D2A was cloned from our own resources^24,26^, according to the sequences listed in Sup. Tab. 1.

The CD8 hinge + CD8TM sequence (FVPVFLPAKPTTTPAPRPPTPAPTIASQPLSLRPEACRPAAGGAVHTRGLDFACDIYIWAPLAGTCGVLLLSLVITLYCNHRNR)^16^ was a kind gift from Dr. Sébastien Wälchli, and it exchanged the CD28 hinge + CD28TM sequence IEVMYPPPYLDNEKSNGTIIHVKGKHLCPSPLFPGPSKPFWVLVVVGGVLACYSLLVTVAFIIFWV.

mEOS4b was cloned from Addgene #54814. tCD19-mCherry was assembled from human CD19 sequence (a kind gift from Prof. Hans-Christian Åsheim^39^), truncation was done according to Addgene #174610^40^, with mCherry sequence cloned from pcDNA3 construct of our own resources. All constructs were verified by Sanger sequencing (GATC Eurofins).

### Lentiviral transduction

Lentiviral particles were produced via PEI or calcium phosphate transfection of Lentix cells. In short, 3-4 mln Lentix cells were seeded a day before on 10cm diameter culture plates. In all cases 15µg of plasmids were used per plate (7,5µg of transfer plasmid, 3,75µg of VSV-G coding plasmid, and 3,75µg of Gag, Pol, Rev, Tat coding plasmid (a kind gift from prof. Johannes Huppa)). PEI transfection was conducted at 1:14 ratio, i.e. 210µg of PEI (MW 25kDa) (Polysciences) to 15µg of DNA. Media was exchanged after 4h. Calcium phosphate transfection was conducted with 0,155M CaCl_2_ (final) in Hepes buffered saline (HBS) buffer with pH 7,05. Media was exchanged after 24h. In both methods lentivirus-containing media was collected 72h after transfection, filtered with 0,45µm filters and immediately added to transduced cells with polybrene at 5ug/ml (final). Human peripheral T-cells were activated a day before with Human T-activator CD3/CD28 Dynabeads (Thermofisher), which were removed prior to transduction. Cells were transduced through spinfection (90min/1200g/32°C) at the concentration 0,25 mln per 1 ml of lentiviral-containing media on 24-wells plates. After the spinning the media was exchanged for the most optimal for given cells. Stably transduced cells were expanded and used for experiments or enriched. Enrichment was achieved either through Dynabeads CD34 positive isolation kit (Thermofisher) or with fluorescence-activated cell sorting (BD FACSMelody Cell Sorter).

### CAR surface expression analysis

Transduced primary T cells were collected, washed and resuspended in PBS with 2%BSA and 2mM EDTA. Cells were stained for 20 min at room temperature (25°C) with SuperFolder CD19-AF647^17^, anti-CD34-FITC (eBioscience), and anti-PD1-PE (Immunotools). Afterwards, they were washed again and analyzed with Sony ID 7000 spectral analyzer.

### Immunofluorescence (IF)

Cells were allowed to adhere to 8-well, glass-bottomed Lab-Tek Chamber Slides (ThermoFisher) in PBS for 45 min at 37°C and 5% CO2, as described elsewhere^41^, followed by fixation with 4% paraformaldehyde (PFA, Merck) in PBS at room temperature (RT) for 12 min. Cells were washed twice gently with PBS and permeabilized with 0.1% Triton X-100 (Merck) in PBS for 15 min at RT, followed by two gentle washes with PBS. Cells were incubated with 1% bovine serum albumin (BSA, Bio-Rad) in PBS for 1 hr at RT. Plastic chambers were then removed, and cells were incubated with primary unconjugated antibody diluted in 1% BSA in PBS for 1 hr at RT. Following primary staining, cells were washed twice gently with 1% BSA in PBS. For secondary staining for IF, cells were incubated for 1 hr in the dark at RT with fluorophore-conjugated secondary antibodies. Cells were counterstained with DAPI and AF546-conjugated phalloidin, in 1% BSA in PBS for 15 min in the dark at RT, washed twice with 1% BSA in PBS, before being mounted with SlowFade Gold (ThermoFisher) antifade reagent. Mounted slides were then sealed with nail polish, and fluorescent signals allowed to mature at –20°C overnight before imaging by confocal microscopy.

### Tetramerized peptides

1ng of SH2D2A biotinylated peptides (SH2D2A Tyr^290^ (Biotin-QPIAFTyrAMG-NH2), SH2D2A pTyr^290^ (Biotin-QPIAFpTyrAMG-NH2), SH2D2A Tyr^305^ (Biotin-APSNITyrVEVED-NH2), SH2D2A pTyr^305^ (Biotin-APSNIpTyrVEVED-NH2)^24^ (GeneCust, Dudelange, Luxembourg) were conjugated with SA-APC (Agilent Technologies PJ27S) in 1:6 molar ratio for at least 17h at 4°C. 6,5ng of the tetramer were used per staining.

### Proximity ligation assay (PLA)

The PLA was performed similarly to IF, based on protocols described elsewhere^42,43^, with cells instead permeabilized with 0.5% Triton X-100 in PBS for 15 min at room temperature (25°C). Primary antibody incubation for PLA occurred overnight at 4°C in a humidity chamber. PLA was accomplished using the following Merck Duolink® reagents: In Situ PLA® Probe Anti-Mouse MINUS, Anti-Rabbit PLUS, and In Situ Detection Reagents FarRed. The PLA reaction was carried out as per the manufacturer’s instructions. Slides were then counterstained with DAPI and phalloidin and mounted as detailed above for the procedure for immunofluorescence.

### Confocal microscopy and image processing

Microscopy slides were imaged on a Zeiss LSM 710 confocal microscope at 63x magnification. Captured images were pre-processed in Zen (Carl Zeiss Imaging) before being loaded into Fiji^44^. Representative images were then selected for each image, and Fiji was used to split and export each colour channel as a separate image.

### Lck SH2 domain GST fusion proteins

Plasmids encoding Lck-SH2 domains WT, Y192F or Y192E GST fusion proteins were expressed in BL21 Codon plus bacteria (Stratagene) and purified on glutathione Sepharose beads (Amersham Biosciences) as described previously^24^. Protein purity and concentrations were assessed with SDS-PAGE and Coomassie Brilliant Blue staining, and by the BCA assay (ThermoFisher) respectively. GST tagged Lck-SH2 domain proteins were cut with 1 unit of PreScission Protease (GE) per 100µg of bound protein in MOPS buffer (50mM MOPS, 50mM NaCl, 1mM DTT, pH 6.8), leaving only the Lck SH2 domains. Purified protein was run on a non-reducing 15% SDS-PAGE gel and stained with Coomassie blue to check protein purity and the lack of protein dimers. Lck SH2 domains were used in affinity measurements by surface plasmon resonance (SPR).

### Surface plasmon resonance (SPR)

The binding profiles of phosphorylated/unphosphorylated peptides to purified WT, Y192E or Y192F Lck-SH2 domains were assessed by SPR analysis on a BIAcore T200 instrument (GE). All SPR analysis were performed at 25°C in HBS-EP buffer (GE, 10mM HEPES pH 7.4, 150mM NaCl, 3mM EDTA, 0.005% v/v Surfactant P20). Biotinylated peptides were immobilized on a streptavidin-coated SA sensor chip (GE) by injection, until a constant level of response units (80-100 RU) was obtained. Varying concentrations of Lck-SH2 domains were passed over the immobilized peptides at a flow rate of 5ul/min, 30s contact time and 100s dissociation time. To remove the Lck-SH2 domains from the peptide, the chip was regenerated with 50mM NaOH at a flow rate of 30μl/min, contact time 30s and 30s stabilization period before the next injection.

### siRNA-mediated knockdown of Lck

siRNA against the human Lck sequence p232 5’-CTG CAA GAC AAC CTG GTT ATC-3’, antisense 5’-TAA CCA GGT TGT CTT GCA GTG-3’ (Eurogentec S.A.), were previously described^45^. Mutated siRNA directed towards the human Lck sense p232 5’-GTG CAA CAC AAC GTG GTT ATC-3’ and antisense 5’-TAA CCA CGT TGT GTT GCA CTG-3’ (Eurogentec S.A.), were used as negative controls. Cells were transfected by electroporation with a BTX electroporator (Genetronix). 15×10^6^ cells and ∼5-µg siRNA were resuspended in 400 µl RPMI-1640 medium without antibiotics supplemented with 5% FCS and transfected at 240V for 25 ms. After 6h, cells were stimulated with PMA/IO.

### Targeted mutagenesis with CRISPR/Cas9

The full protocol and methodological considerations of generation of the Jurkat T cells knock-in mutants are described elsewhere^23^. Jurkat T cells (10 – 15 mln) were co-transfected with 5 µg Cas9 coding DNA plasmid (pX330-U6-Chimeric-BB-CBh-hSpCas9, Addgene) with guiding sequences designed to target *SH2D2A* in exon 7 at the codon for Tyr^290^, 0,5 µg pEGFP-N1 plasmid (Clontech), and 1µM single stranded DNA repair sequences (SH2D2A Y290F and Y290E) by electroporation with a BTX electroporator (Genetronix) as described above. After three days, transfected cells were cloned by limiting dilution. Three weeks later clones were screened with allele-specific PCR. Generated mutations were confirmed with sequencing (GATC Biotech). SH2D2A KO cell lines were obtained as byproducts of this procedure.

### Stimulation of cells

PMA/IO stimulation: cells were stimulated with 50 ng/ml PMA and 500 ng/ml IO for 17h. Cells were then washed and rested for 24h. Phosphatase inhibition by PV (pervanadate) treatment: up to 10^7^ cells/ml in PBS were prewarmed at 37°C water bath for 5 min. Cells were treated with 0,01mM sodium orthovanadate (Sigma) and 0,01% hydrogen peroxide (Sigma-Aldrich) for 5 min. Treatment was stopped by sample transfer to 4°C PBS. Inhibition of Src kinases by PP2 treatment: cells were treated with 10 µM PP2 for 45 min at 37°C. Short-term TCR stimulation: up to 10^8^ cells/ml in PBS were prewarmed at 37°C for 5 min. Cells were stimulated with 5µg/ml anti-TCR antibody (OKT3 or C305), and if specified, with anti-CD28 antibody, for 2 min. Stimulation was stopped by adding 1 ml of cold PBS and cells were collected for lysis. Long-term TCR stimulation: cells were cultured on plates or flasks previously coated with 5µg/ml of anti-CD3 (OKT3) in PBS and incubated for 17h at 37°C. Cells were then washed and rested in new plates/flasks for 24h.

### Lysis, immunoprecipitation, pull-down and western blotting

Stimulated or treated cells were pelleted and lysed with 0.1% LDS/ 1% Triton solution containing 0,5% Triton X-100 (VWR international S.A.S), 50mM HEPES (ThermoFisher), 0,05% LDS (Merck), 0.05 M LiCl, 0.5 mM PMSF, 2.5 mM EDTA (pH 8.0), 1mM sodium vanadate and 1x SIGMAFAST protease inhibitor cocktail (all from Sigma). Cells were lysed for 45 min on ice. Lysates were sonicated briefly to break the DNA. Prior to immunoprecipitation (IP), lysates were preabsorbed using protein G Dynabeads (ThermoFisher). Protein G Dynabeads were incubated with anti-SH2D2A antibody at room temperature for 1 hour. Antibody-treated beads were washed and then incubated with lysates on a rotating wheel at 4°C for 1 hour. Myone Streptavidin T1 Dynabeads (ThermoFisher) covered by biotinylated SH2D2A peptides were used analogically for SH2D2A peptides pull-down (PD). After IP, beads were washed three times with lysis buffer. The appropriate volume of loading buffer (containing 0.35 M Tris HCl, 10% SDS, 6% β-mercaptoethanol (all Sigma), 30% glycerol (VWR international S.A.S), 0,175mM bromophenol blue (Fluka Ag), pH 6.8) was added to the lysates and the IP beads. All samples were denatured by boiling at 95°C for 10 min. Denatured samples were run with SDS-PAGE and transferred onto PVDF membranes (Bio-Rad laboratories) with Trans-Blot Turbo Transfer System (Bio-Rad laboratories). The membranes were incubated with primary antibodies diluted in tris-buffered saline (pH 7.4), 0,1% Tween (Sigma-Aldrich), and 3% skimmed milk or 3% bovine serum albumin (Bio-Rad laboratories) (when combined with anti-pTyr antibodies). After overnight incubation at 4°C, the membranes were incubated with appropriate HRP-conjugated secondary antibody for 1h at room temperature. SuperSignal West Pico Stable Peroxide Solution (Pierce) was used to visualize bands using ChemiDoc^TM^ Imaging System (Bio-Rad Laboratories). Images were quantified with ImageJ software. Normalization method for each experiment is given in the results section or in figures legends.

### Mass spectrometry analysis

Samples for label-free quantitative proteomics were prepared according to the protocol for SH2D2A peptides PD. Afterwards, proteins were reduced, alkylated and digested directly on beads with ProteaseMAX™ Surfactant, 3,6 μg of trypsin (Sequencing grade modified, Promega) overnight at 37°C, according to the manufacturer’s protocol. Samples were desalted with STAGE-TIP method^46^ using C18 resin disk (3M Empore). Peptide elution was performed with 0,1% formic acid and 80% acetonitrile. Eluates were concentrated with SpeedVac until a volume of approximately 7 µl. All experiments were performed on an Easy nLC1000 nano-LC system connected to a quadrupole – Orbitrap (QExactive) mass spectrometer (ThermoElectron, Bremen, Germany) equipped with a nanoelectrospray ion source (EasySpray/Thermo). Liquid chromatography separation was performed with an EasySpray column (C18, 2 µm beads, 100 Å, 75 μm inner diameter) (Thermo) capillary of 25 cm bed length. The flow rate used was 300 nL/min, and the solvent gradient was 2 % B to 30 % B in 120 minutes, then 90 % B wash in 20 minutes. Solvent A was aqueous 0.1 % formic acid, whereas solvent B was 100 % acetonitrile in 0.1 % formic acid. Column temperature was kept at 60°C. The mass spectrometer was operated in the data-dependent mode to automatically switch between MS and MS/MS acquisition. Survey full scan MS spectra (from m/z 400 to 1,200) were acquired in the Orbitrap with resolution R = 70,000 at m/z 200 (after accumulation to a target of 3,000,000 ions in the quadruple). The method used allowed sequential isolation of the most intense multiply-charged ions, up to ten, depending on signal intensity, for fragmentation on the HCD cell using high-energy collision dissociation at a target value of 100,000 charges or maximum acquisition time of 100 ms. MS/MS scans were collected at 17,500 resolution at the Orbitrap cell. Target ions already selected for MS/MS were dynamically excluded for 30 seconds. General mass spectrometry conditions were: electrospray voltage, 2.1 kV; no sheath and auxiliary gas flow, heated capillary temperature of 250°C, normalized HCD collision energy 25%. Ion selection threshold was set to 1e4 counts. Isolation width was 3.0 Da. MS raw files were submitted to MaxQuant software version 1.4.0.8^47^ for protein identification. Parameters were set as follow: protein N-acetylation, methionine oxidation and pyroglutamate conversion of Glu and Gln as variable modifications. First search error window of 20 ppm and main search error window of 6 ppm. Trypsin without proline restriction enzyme option was used with two miscleavages allowed. Minimal unique peptides were set to 1, and FDR was 0.01 (1%) for peptide and protein identification. Label-free quantitation was set with a retention time alignment window of 3 min. The Uniprot human database was used (December 2013). Generation of reversed sequences was selected to assign FDR rates. The MS data will be available in the PRIDE database^48^ under accession number XXX after the manuscript has been accepted for publication.

### CAR T-cell: target cell co-culture experiment

Not-enriched, stably transduced primary human CD8^+^ CAR T-cells were seeded with stably transduced (mEOS4b or tCD19-mCherry) cancer cell lines (THP1, WM35, OVCAR3, and SKBR3) at ratio 1:1 (200k cells: 200k cells) on 48-wells plates in 200ul of primary T-cells media supplemented with 100U/ml IL2. Prior to seeding, CAR T-cells were treated with CellTracker Blue CMAC (Invitrogen) according to manufacturer’s protocol.

Samples were collected after 48h (2 days) and 168h (7 days). Cells incubated for 7 days were fed with extra media (supplemented with 200U/ml IL2) on day 2 and 5. At given timepoints cells were detached and collected with FACS buffer (1%BSA, 2mM EDTA in PBS) and divided into two samples, for intracellular and extracellular staining.

Samples were first treated with LIVE/DEAD Fixable Blue Dead Cell Stain (Invitrogen) according to manufacturer’s protocol, followed by Fc Block (eBioscience™, Invitrogen) in FACS buffer for 20 min at RT. Prior to live/dead stain and Fc Block, samples for intracellular staining were incubated with Brefeldin A (eBioscience™, Invitrogen) in T cells media at 37°C for 4hrs. After Fc Block, samples for intracellular staining were fixed and permeabilized with eBioscience™ Foxp3/ Transcription Factor Staining Buffers (Invitrogen) according to manufacturer’s protocol.

The choice of markers for the panel is explained in Sup. Tab. 2. Extracellular staining was started with antibodies against chemokine receptors (CCR7 (CD197)-PrCPE Fluor 710 (Thermofisher); CCR6 (CD196)-BV711, CXCR3 (CD183)-PE Fire 640, CCR4 (CD194)-PECF594 (Sony)) diluted in Brilliant Stain Buffer (Invitrogen) and incubated with samples for 10 min at RT. It was continued with addition of antibodies against other surface markers (CD45RA-Nova Fluor Blue 660-120S, CD8a-BUV805, TIM3 (CD366)-PE Cy 5.5, CD56 (NCAM)-BUV737, CD69-BUV395, CD27-BUV661, CD4-Nova Fluor Blue 585, CD28-BUV615 (Thermofisher), CD45RO-BV510, CD3-AF700, CD127 (IL7R)-APC Cy7, CD25-BV605 or CD25-PE (Immunotools), PD-1 (CD279)-BV421 (Sony) or PD1-R718 (BD Biosciences), CD34-APC (eBioscience), CD107a-V450 (BD Biosciences)) diluted in CellBlox Buffer (Invitrogen) and FACS buffer and incubated with samples for 20 min at RT.

Intracellular staining was done with antibodies against intracellular and surface markers (Granzyme B-PE Cy 7, IL-2-PE, FOXP3-BV785, CD25-BV605, PD-1 (CD279)-BV421, TNFα-BV650 (Sony), CD8a-BUV805, TIM3-PE Cy5.5, IFNγ-BUV496 (Thermofisher), CD34-APC (eBioscience), CD107a-V450 (BD Biosciences), CD25-NovaFluorRed700 (Thermofisher)) diluted in CellBlox Buffer (Invitrogen) and FACS buffer and incubated with samples for 72h at 4°C.

All samples were washed three times before being analysed with Sony ID 7000 spectral analyzer. Time-wise matched samples (2^nd^ and 7^th^ day) were run with the same voltage settings.

### CAR T-cell: target cell co-culture analysis

#### Statistical analysis

GraphPad Prism version 7.04 was used for statistical analysis, except of MS data, which was analyzed with Perseus version 1.5.8.5. Statistical significance of the results was assessed with unpaired, two tailed t-tests. All statistically significant comparisons are shown on the graphs accordingly: * - p-value <0.05; ** - p-value <0.01; *** - p-value <0.001.

## Author contributions

P.B. and A.S. conceptualized the study. P.B., H.C., B.C.G., R.P.G., V.S. and A.S. designed experiments and analyzed results. P.B. performed FACS, immunoprecipitation, pulldowns, immunoblotting, lentiviral transductions, CRISPR/Cas9, co-culture assays, immunofluorescence and flow cytometry. B.C.G. performed immunofluorescence, microscopy, proximity ligation assay, and co-culture assays data analysis. H.C. performed FACS, lentiviral transductions, and flow cytometry. R.P.G. performed SH2D2A peptides pulldown. G.A.S. performed mass spectrometry. P.B. and V.S. performed western blotting. H.C. and S.F. performed surface plasmon resonance. H.C. produced recombinant LCK-SH2 proteins. J.S. and M.T. designed and produced CD19 SuperFolder. T.P. and R.P. provided training and experimental assistance. P.B., B.C.G., and J.G.V. performed PBMC isolation. P.B., H.C., J.G.V., and A.P. performed co-culture assays. P.B. designed plasmids. P.B., H.K., M.D., I.A., and S.L.A. made plasmids. P.B. and A.S. performed sequence conservation analysis. S.W. and J.H. assisted with experimental design and reagents. P.B. wrote the manuscript with input from all co-authors. A.S. oversaw the project and reviewed the final manuscript.

## Acknowledgements

The study was support by the Norwegian Cancer Society (grant numbers 107561 and 208360), the Norwegian Research Council (grant numbers 214202 and 302647), Stiftelsen Anyes, Anders Jahres foundation, Unifor, the University of Oslo, Nansenfondet og de dermed forbundne fond, Astri og Birger Torsteds legat til bekjempelse av kreft, Familien Blix’ Fond Til Fremme Av Medisinsk Forskning, EMBO Scientific Exchange Grant Number 8984. Schematic figures were created with BioRender.

## References

1. June, C. H., O’Connor, R. S., Kawalekar, O. U., Ghassemi, S. & Milone, M. C. CAR T cell immunotherapy for human cancer. Science 359, 1361–1365 (2018).

2. Bouchkouj, N. et al. FDA Approval Summary: Axicabtagene Ciloleucel for Relapsed or Refractory Large B-cell Lymphoma. Clin. Cancer Res. 25, 1702–1708 (2019).

3. Sharma, P. et al. FDA Approval Summary: Idecabtagene Vicleucel for Relapsed or Refractory Multiple Myeloma. Clin. Cancer Res. 28, 1759–1764 (2022).

4. Gudipati, V. et al. Inefficient CAR-proximal signaling blunts antigen sensitivity. Nat. Immunol. 21, 848–856 (2020).

5. Demetriou, P. et al. A dynamic CD2-rich compartment at the outer edge of the immunological synapse boosts and integrates signals. Nat. Immunol. 21, 1232–1243 (2020).

6. Dong, R. et al. Rewired signaling network in T cells expressing the chimeric antigen receptor (CAR). EMBO J. 39, (2020).

7. Stoiber, S. et al. Limitations in the Design of Chimeric Antigen Receptors for Cancer Therapy. Cells 8, 472 (2019).

8. Wu, L. et al. CD28-CAR-T cell activation through FYN kinase signaling rather than LCK enhances therapeutic performance. Cell Rep. Med. 4, 100917 (2023).

9. Woessner, N. M., Uleri, V., Stepanek, O. & Minguet, S. The TCR and LCK: foundations for T-cell activation and therapeutic innovation. Front. Immunol. 16, 1737013 (2026).

10. Geiger, T. L., Nguyen, P., Leitenberg, D. & Flavell, R. A. Integrated src kinase and costimulatory activity enhances signal transduction through single-chain chimeric receptors in T lymphocytes. Blood 98, 2364–2371 (2001).

11. Hartl, F. A. et al. Noncanonical binding of Lck to CD3ε promotes TCR signaling and CAR function. Nat. Immunol. 21, 902–913 (2020).

12. Suryadevara, C. M. et al. Preventing Lck Activation in CAR T Cells Confers Treg Resistance but Requires 4-1BB Signaling for Them to Persist and Treat Solid Tumors in Nonlymphodepleted Hosts. Clin. Cancer Res. 25, 358–368 (2019).

13. Borowicz, P., Chan, H., Hauge, A. & Spurkland, A. Adaptor proteins: Flexible and dynamic modulators of immune cell signalling. Scand. J. Immunol. 92, e12951 (2020).

14. Krawczyk, M. et al. The costimulatory domain influences CD19 CAR-T cell resistance development in B-cell malignancies. Preprint at 10.1101/2025.02.28.640707 (2025).

15. Norell, H. et al. CD34-based enrichment of genetically engineered human T cells for clinical use results in dramatically enhanced tumor targeting. Cancer Immunol. Immunother. 59, 851–862 (2010).

16. Bajor, M. et al. PD-L1 CAR effector cells induce self-amplifying cytotoxic effects against target cells. J. Immunother. Cancer 10, e002500 (2022).

17. Laurent, E. et al. Directed Evolution of Stabilized Monomeric CD19 for Monovalent CAR Interaction Studies and Monitoring of CAR-T Cell Patients. ACS Synth. Biol. 10, 1184–1198 (2021).

18. Xu, X. et al. Phase separation of chimeric antigen receptor promotes immunological synapse maturation and persistent cytotoxicity. Immunity 57, 2755–2771.e8 (2024).

19. Wang, P. et al. Chimeric antigen receptor with novel intracellular modules improves antitumor performance of T cells. Signal Transduct. Target. Ther. 10, (2025).

20. Velasco Cárdenas, R. M.-H., et al. Harnessing CD3 diversity to optimize CAR T cells. Nat. Immunol. 24, 2135–2149 (2023).

21. Feucht, J. et al. Calibration of CAR activation potential directs alternative T cell fates and therapeutic potency. Nat. Med. 25, 82–88 (2019).

22. Granum, S. et al. Modulation of Lck Function through Multisite Docking to T Cell-specific Adapter Protein. J. Biol. Chem. 283, 21909–21919 (2008).

23. Borowicz, P. et al. A simple and efficient workflow for generation of knock-in mutations in Jurkat T cells using CRISPR/Cas9. Scand. J. Immunol. 91, e12862 (2020).

24. Granum, S. et al. The kinase Itk and the adaptor TSAd change the specificity of the kinase Lck in T cells by promoting the phosphorylation of Tyr ^192^. Sci. Signal. 7, (2014).

25. Pelosi, M. et al. Tyrosine 319 in the Interdomain B of ZAP-70 Is a Binding Site for the Src Homology 2 Domain of Lck. J. Biol. Chem. 274, 14229–14237 (1999).

26. Spurkland, A. et al. Molecular Cloning of a T Cell-specific Adapter Protein (TSAd) Containing an Src Homology (SH) 2 Domain and Putative SH3 and Phosphotyrosine Binding Sites. J. Biol. Chem. 273, 4539–4546 (1998).

27. Hinckley-Boned, A. et al. Tailoring CAR surface density and dynamics to improve CAR-T cell therapy. J. Immunother. Cancer 13, e010702 (2025).

28. Balagopalan, L. et al. Generation of antitumor chimeric antigen receptors incorporating T cell signaling motifs. Sci. Signal. 17, eadp8569 (2024).

29. Cheng, B., Montmasson, M., Terradot, L. & Rousselle, P. Syndecans as Cell Surface Receptors in Cancer Biology. A Focus on their Interaction with PDZ Domain Proteins. Front. Pharmacol. 7, (2016).

30. Bonifacino, J. S. & Dell’Angelica, E. C. Molecular Bases for the Recognition of Tyrosine-based Sorting Signals. J. Cell Biol. 145, 923–926 (1999).

31. Cai, C. Y., Zhang, X., Sinko, P. J., Burakoff, S. J. & Jin, Y.-J. Two Sorting Motifs, a Ubiquitination Motif and a Tyrosine Motif, Are Involved in HIV-1 and Simian Immunodeficiency Virus Nef-Mediated Receptor Endocytosis. J. Immunol. 186, 5807–5814 (2011).

32. Barbera, S., et al. Trogocytosis of chimeric antigen receptors between T cells is regulated by their transmembrane domains. Sci. Immunol. 10, eado2054 (2025).

33. Gioia, L., Siddique, A., Head, S. R., Salomon, D. R. & Su, A. I. A genome-wide survey of mutations in the Jurkat cell line. BMC Genomics 19, (2018).

34. Schneider, U., Schwenk, H. & Bornkamm, G. Characterization of EBV-genome negative “null” and “T” cell lines derived from children with acute lymphoblastic leukemia and leukemic transformed non-Hodgkin lymphoma. Int. J. Cancer 19, 621–626 (1977).

35. Clipstone, N. A. & Crabtree, G. R. Identification of calcineurin as a key signalling enzyme in T-lymphocyte activation. Nature 357, 695–697 (1992).

36. Thiruchelvam-Kyle, L. et al. The Activating Human NK Cell Receptor KIR2DS2 Recognizes a β2-Microglobulin–Independent Ligand on Cancer Cells. J. Immunol. 198, 2556–2567 (2017).

37. Gibson, D. G. et al. Enzymatic assembly of DNA molecules up to several hundred kilobases. Nat. Methods 6, 343–345 (2009).

38. Forloni, M., Liu, A. Y. & Wajapeyee, N. Megaprimer Polymerase Chain Reaction (PCR)-Based Mutagenesis. Cold Spring Harb. Protoc. 2019, pdb.prot097824 (2019).

39. Stamenkovic, I. & Seed, B. CD19, the earliest differentiation antigen of the B cell lineage, bears three extracellular immunoglobulin-like domains and an Epstein-Barr virus-related cytoplasmic tail. J. Exp. Med. 168, 1205–1210 (1988).

40. Veatch, J. R. et al. A therapeutic cancer vaccine delivers antigens and adjuvants to lymphoid tissues using genetically modified T cells. J. Clin. Invest. 131, (2021).

41. Tsang, M., Gantchev, J., Ghazawi, F. M. & Litvinov, I. V. Protocol for Adhesion and Immunostaining of Lymphocytes and Other Non-Adherent Cells in Culture. BioTechniques 63, 230–233 (2017).

42. Alam, M. S. Proximity Ligation Assay (PLA). Curr. Protoc. Immunol. 123, (2018).

43. Hegazy, M. et al. Proximity Ligation Assay for Detecting Protein-Protein Interactions and Protein Modifications in Cells and Tissues in Situ. Curr. Protoc. Cell Biol. 89, (2020).

44. Schindelin, J., et al. Fiji: an open-source platform for biological-image analysis. Nat. Methods 9, 676–682 (2012).

45. Methi, T. et al. Short-Interfering RNA-Mediated Lck Knockdown Results in Augmented Downstream T Cell Responses. J. Immunol. 175, 7398–7406 (2005).

46. Rappsilber, J., Ishihama, Y. & Mann, M. Stop and Go Extraction Tips for Matrix-Assisted Laser Desorption/Ionization, Nanoelectrospray, and LC/MS Sample Pretreatment in Proteomics. Anal. Chem. 75, 663–670 (2003).

47. Cox, J. & Mann, M. MaxQuant enables high peptide identification rates, individualized p.p.b.-range mass accuracies and proteome-wide protein quantification. Nat. Biotechnol. 26, 1367–1372 (2008).

48. Perez-Riverol, Y. et al. The PRIDE database and related tools and resources in 2019: improving support for quantification data. Nucleic Acids Res. 47, D442–D450 (2019).

49. Némorin, J.-G. & Duplay, P. Evidence That Lck-mediated Phosphorylation of p56 and p62 May Play a Role in CD2 Signaling. J. Biol. Chem. 275, 14590–14597 (2000).

50. Jung, S. H. et al. ARAP, a Novel Adaptor Protein, Is Required for TCR Signaling and Integrin-Mediated Adhesion. J. Immunol. 197, 942–952 (2016).

51. Shapiro, M. J., Nguyen, C. T., Aghajanian, H., Zhang, W. & Shapiro, V. S. Negative Regulation of TCR Signaling by Linker for Activation of X Cells via Phosphotyrosine-Dependent and -Independent Mechanisms. J. Immunol. 181, 7055–7061 (2008).

52. Lo, W.-L. et al. Lck promotes Zap70-dependent LAT phosphorylation by bridging Zap70 to LAT. Nat. Immunol. 19, 733–741 (2018).

53. Kabouridis, P. S., Isenberg, D. A. & Jury, E. C. A negatively charged domain of LAT mediates its interaction with the active form of Lck. Mol. Membr. Biol. 28, 487–494 (2011).

54. Brdička, T. et al. Non–T Cell Activation Linker (NTAL). J. Exp. Med. 196, 1617–1626 (2002).

55. Brdičková, N. et al. LIME. J. Exp. Med. 198, 1453–1462 (2003).

56. Hur, E. M. et al. LIME, a Novel Transmembrane Adaptor Protein, Associates with p56lck and Mediates T Cell Activation. J. Exp. Med. 198, 1463–1473 (2003).

57. Brdci ka, T. et al. Phosphoprotein Associated with Glycosphingolipid-Enriched Microdomains (Pag), a Novel Ubiquitously Expressed Transmembrane Adaptor Protein, Binds the Protein Tyrosine Kinase Csk and Is Involved in Regulation of T Cell Activation. J. Exp. Med. 191, 1591–1604 (2000).

58. Huang, X., Li, Y., Tanaka, K., Moore, K. G. & Hayashi, J. I. Cloning and characterization of Lnk, a signal transduction protein that links T-cell receptor activation signal to phospholipase C gamma 1, Grb2, and phosphatidylinositol 3-kinase. Proc. Natl. Acad. Sci. 92, 11618–11622 (1995).

59. Li, Y., He, X., Schembri-King, J., Jakes, S. & Hayashi, J. Cloning and Characterization of Human Lnk, an Adaptor Protein with Pleckstrin Homology and Src Homology 2 Domains that Can Inhibit T Cell Activation. J. Immunol. 164, 5199–5206 (2000).

60. Fukushima, A. et al. Lck couples Shc to TCR signaling. Cell. Signal. 18, 1182–1189 (2006).

61. Marie-Cardine, A. et al. SHP2-interacting Transmembrane Adaptor Protein (SIT), A Novel Disulfide-linked Dimer Regulating Human T Cell Activation. J. Exp. Med. 189, 1181–1194 (1999).

62. Marie-Cardine, A. et al. Molecular Cloning of SKAP55, a Novel Protein That Associates with the Protein Tyrosine Kinase p59 in Human T-lymphocytes. J. Biol. Chem. 272, 16077–16080 (1997).

63. Kang, H. SH3 domain recognition of a proline-independent tyrosine-based RKxxYxxY motif in immune cell adaptor SKAP55. EMBO J. 19, 2889–2899 (2000).

64. Saitoh, K. et al. STAP-2 Is a Novel Positive Regulator of TCR-Proximal Signals. J. Immunol. 209, 57–68 (2022).

65. Bruyns, E. et al. T Cell Receptor (TCR) Interacting Molecule (TRIM), A Novel Disulfide-linked Dimer Associated with the TCR–CD3–ζ Complex, Recruits Intracellular Signaling Proteins to the Plasma Membrane. J. Exp. Med. 188, 561–575 (1998).

