## Supplementary data for "Intrinsically disordered insert from SH2D2A rewires CD19 CAR signaling via Tyr290"

### Supplementary tables

#### Supplementary table 1

List of Lck adaptor proteins identified through the literature search for the purpose of this study*

| Protein name | Interaction with Lck | Lck-binding motif | Ref. |
|---|---|---|---|
| CD3ε (control) | It is a part of the TCR, necessary for its correct signal transduction. It contains one ITAM motif. The RK-motif in CD3ε recognized by LCK has been successfully applied in CAR design, improving its function. | 153-202aa<br>KNRKAKAKPVTRGAGAGGRQRGQNKERPPPVPNP DYEPI**RKGQRDLY**SGL<br>Lck SH3 domain recognizes a non-canonical motif RKGQRDLY | [11] |
| DOK1 and DOK2 | Upon phosphorylation, DOK1 and DOK2 are binding the Lck-SH2 domain. | Unidentified motif. Requires further investigation. | [49] |
| FYB1 and FYB2 | Both binds Lck. | Unidentified motif. Requires further investigation. | [50] |
| HSH2D | HSH2D promotes phosphorylation of the adaptor LAX1 by Lck, which depends on the HSH2D SH2 domain. No data on binding. | Unidentified motif. Requires further investigation. | [51] |
| LAT | Lck promotes Zap-70 dependent LAT phosphorylation by bridging Zap-70 to LAT.[52] | 71-92aa<br>PPLSQPDLL**PIPRSP**QPLGGSH<br>Lck SH3 domain interacts with the PIPRSP motif.<br>105-129aa (in alternative isoform)<br>NSVASYEN**EEPACEDADEDEDD**YHN<br>Lck interacts with the EEPACEDADEDEDD motif. | [52]<br>[53] |
| LAT2 | It is most phosphorylated in the presence of Lck and ZAP70. No data on binding. | Unidentified motif. Requires further investigation. | [54] |
| LIME1 | Phosphorylated LIME1 binds the Lck SH2 domain. | 226-261aa<br>ILALAGDLA**Y**QTLPLRALDVDSGPLENV**Y**ESIRELG<br>Phosphorylated Tyr235 and Tyr254 are recognized by the Lck SH2 domain | [55,56] |
| PAG1 | PAG1 is phosphorylated by Lck and binds to Lck SH2 domain. | Unidentified motif. Requires further investigation. | [57] |
| SH2B3 | Lck phosphorylates SH2B3. Phosphorylated SH2B3 binds the Lck SH2 domain. | Unidentified motif, but most likely it includes pTyr555. Requires further investigation. | [58,59] |
| SH2D2A | Multivalent interactions with Lck promote SH2D2A phosphorylation. | 236-312aa<br>EKE**PSQLLRPKPPIP**AKPQLPPEVYTIPVPRHRPAPR PKPSNPI**Y**NEPDEPIAF**Y**AMGRGSPGEAPSNI**Y**VEVE DEG<br>Phosphorylated Tyr280, Tyr290, and Tyr305 are recognized by the Lck SH2 domain (especially Tyr305). Proline rich region 239–256aa is recognized by Lck SH3 domain.<br>236-279aa<br>EKE**PSQLLRPKPPIP**AKPQLPPEVYTIPVPRHRPAPR PKPSNPI<br>Only proline rich region recognized by Lck SH3 domain. | [22] |
| SHC1 | SHC1 interacts with LCK SH2/3 domains via its' PTB domain. | Interaction through protein domain and not a motif. | [60] |
| SIT1 | Lck can phosphorylate SIT1. No data on binding. | Unidentified motif. Requires further investigation. | [61] |

| | | | |
|---|---|---|---|
| SKAP1 | SKAP1 binds Lck SH2 domain but is not phosphorylated by Lck. Lck SH3 domain binds 204-299aa, and specifically 277-298aa in SKAP1. | 267-298aa<br>EEDI**Y**EVLPDEEHDLEEDESGTR**RKGVDYASY**Y<br>RKxxYxxY motif is recognized by Lck, extended to include highly conserved Tyr271 motif, possibly recognized by Lck SH2. | [62,63] |
| STAP2 | The STAP-2 PH domain was found to bind to Lck. The interaction required also pTyr250, as Phe250 mutant did not bind Lck. | Interaction through protein domain, however, motif binding might be also involved (potentially Tyr250 and proline-rich regions). Requires further investigation. | [64] |
| TRAT1 | Lck-SH2 domain can pulldown TRAT1 (also known as TRIM), but TRAT1 does not pulldown Lck in T cell lysate. Lck can phosphorylate TRAT1. | Unidentified motif. Requires further investigation. | [65] |

* Gray shading indicates Lck-adaptors and their amino acid sequence motifs incorporated in our CAR design.

#### Supplementary table 2

Definition of markers used in co-culture assay.

| Marker | Purpose |
|---|---|
| Cell tracker blue | Dilution of this marker in T cells correlates with cell division (proliferation) |
| Live/dead | This dye is absorbed only by cells with compromised membrane (dead cells) |
| mEOS4b | Cancer cell lines which do not express CD19 |
| tCD19-mCherry | Cancer cell lines expressing CD19 (CAR target), T cell trogocytosis (recipient) |
| FOXP3 | Treg marker |
| Granzyme B | Cytotoxicity marker |
| IFNg | Proinflammatory cytokine |
| IL-2 | Proinflammatory cytokine, lymphocytes growth factor |
| TNFa | Proinflammatory cytokine |
| CD3 | T cell receptor |
| CD4 | Helper T cells co-receptor |
| CD8a | Cytotoxic T cells co-receptor |
| CD25 | Activation, T regs |
| CD27 | Memory and effector T cells |
| CD28 | T cells, downregulation indicates chronic activation |
| CD34 | CAR T cell transduction |
| CD45RO | Memory T cell |
| CD45RA | Naïve, Stem cell memory T cells (Tscm) |
| CD56 (NCAM) | NK cells, NK-like T cells, binds CD56 |
| CD69 | Activation |
| CD107a | Degranulation |
| CD127 (IL7R) | Downregulation indicates T regs |
| CXCR3 (CD183) | Th1 |
| CCR4 (CD194) | Th2 |
| CCR6 (CD196) | Th17 |
| CCR7 (CD197) | Stem cell memory T cells (Tscm) |
| PD-1 (CD279) | Exhaustion |
| TIM3 (CD366) | Exhaustion |

### Supplementary figures

#### Supplementary figure 1

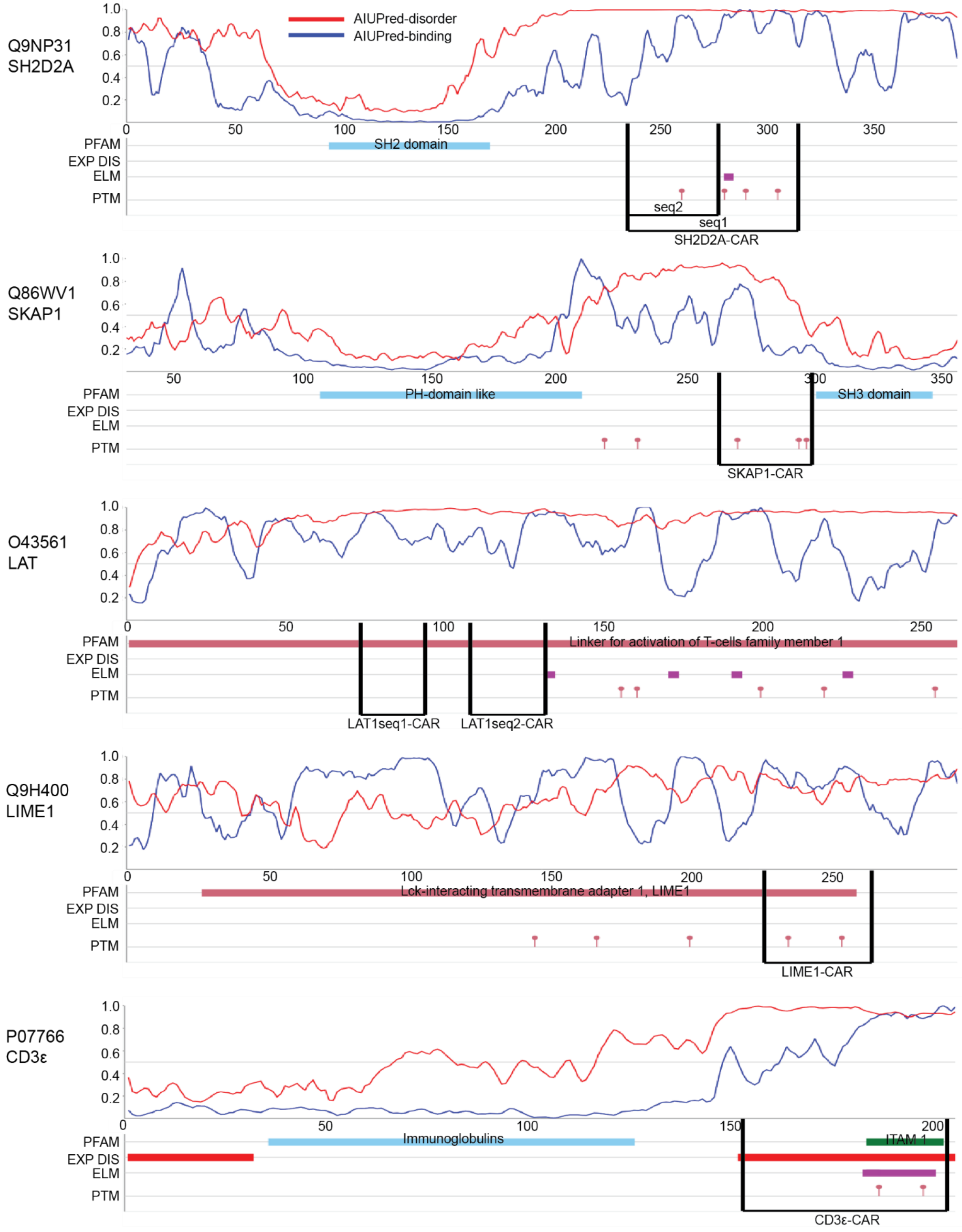

#### Supplementary figure 2

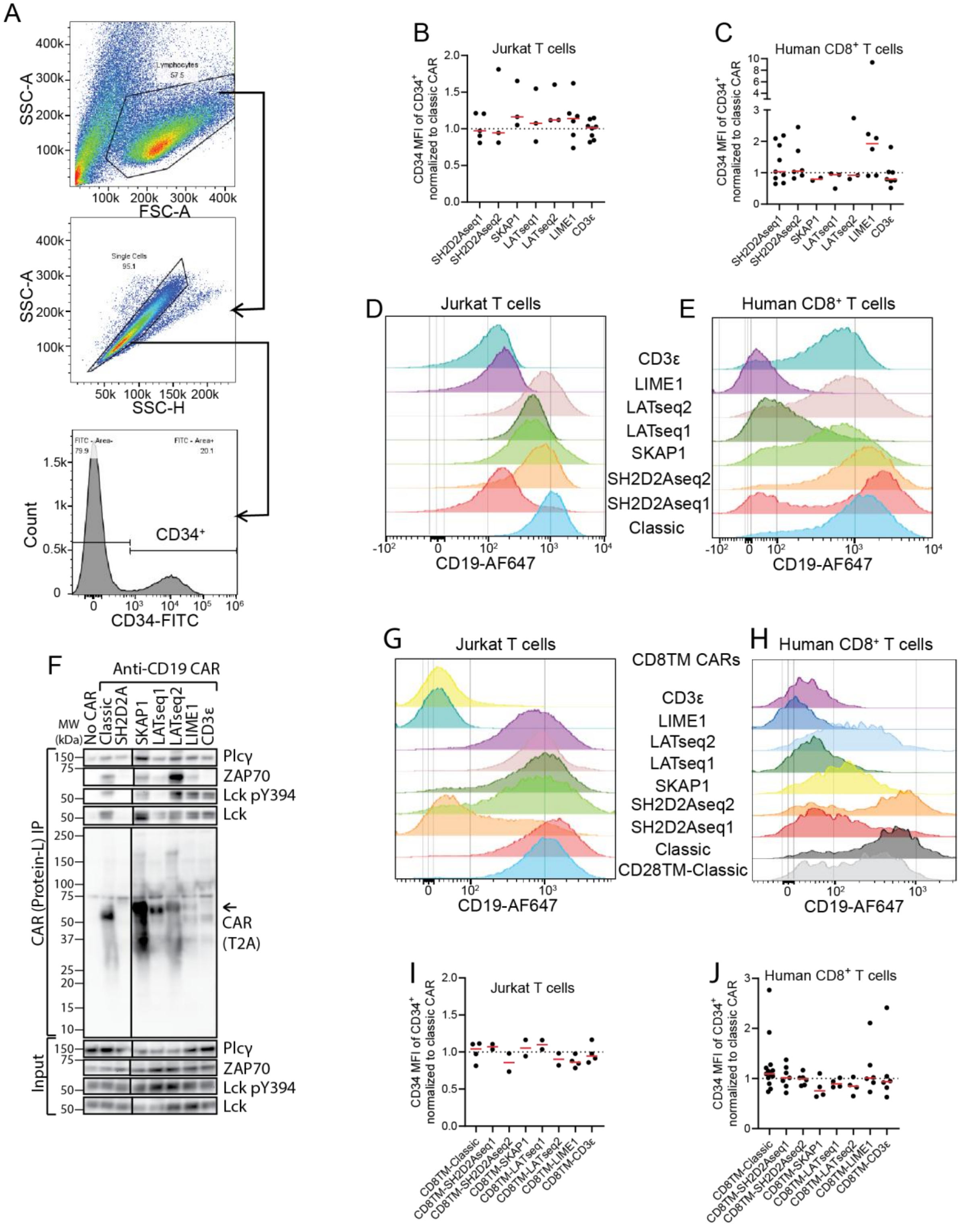

#### Supplementary figure 3

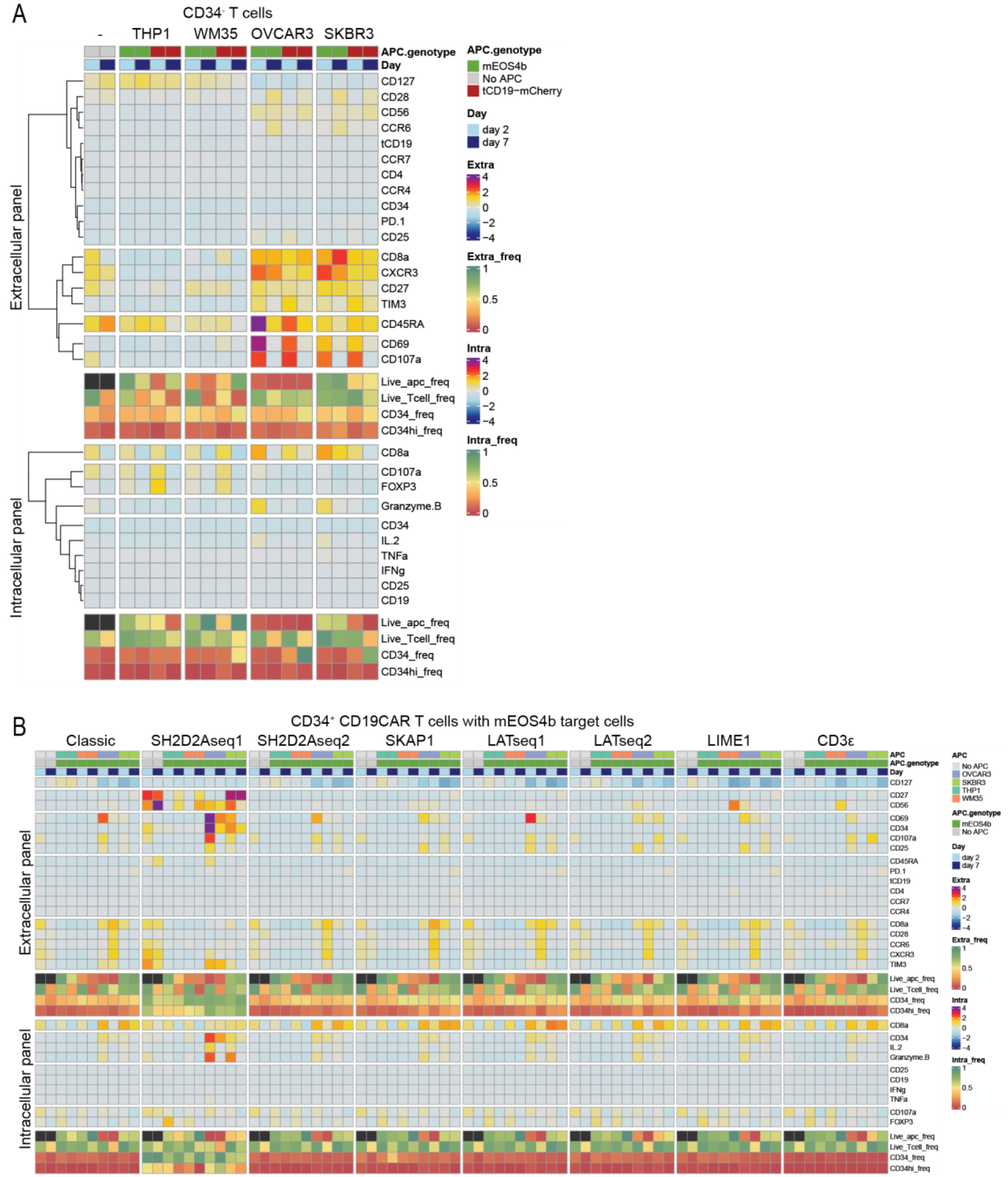

### Supplementary figure 4

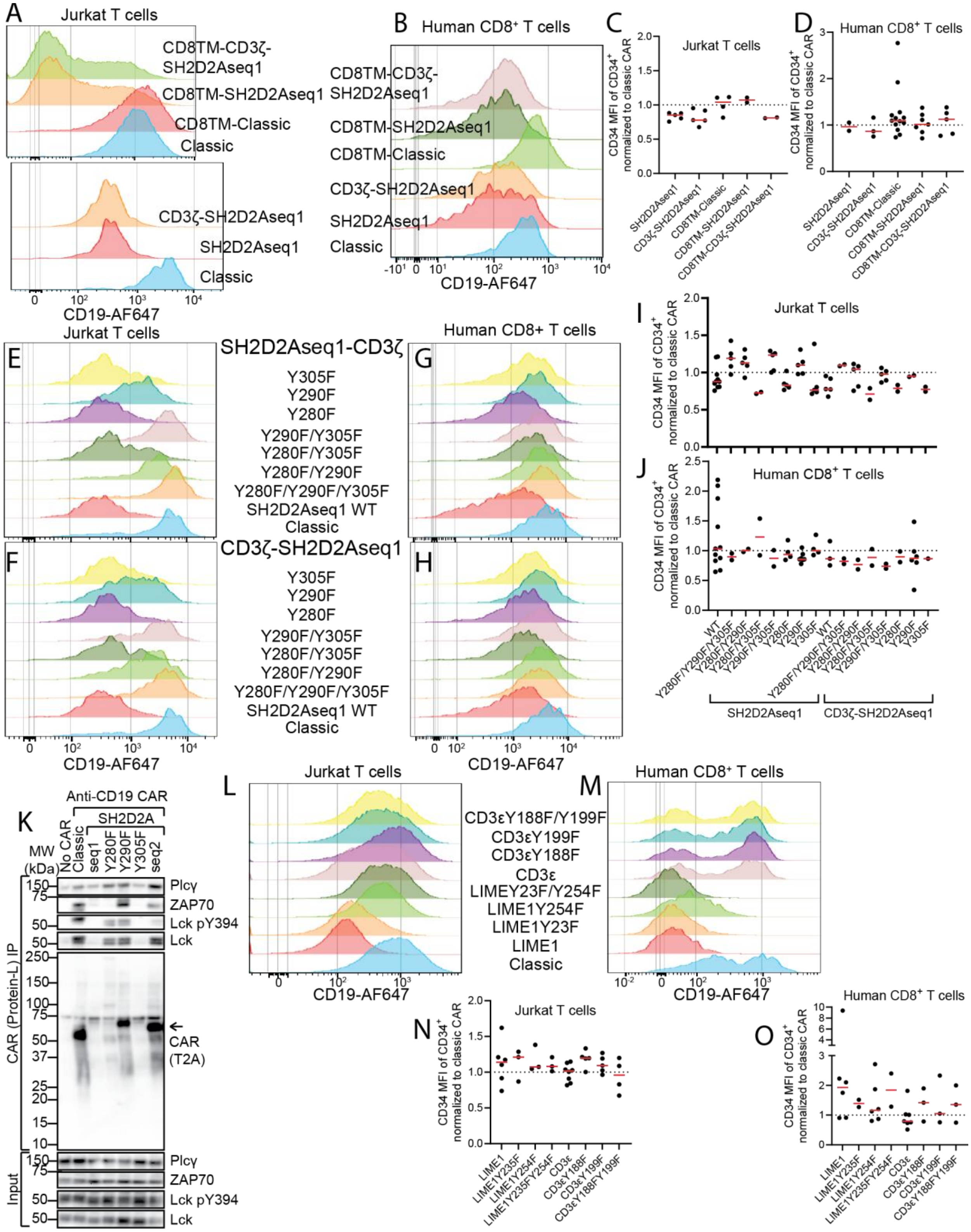

### Supplementary figure 5

A

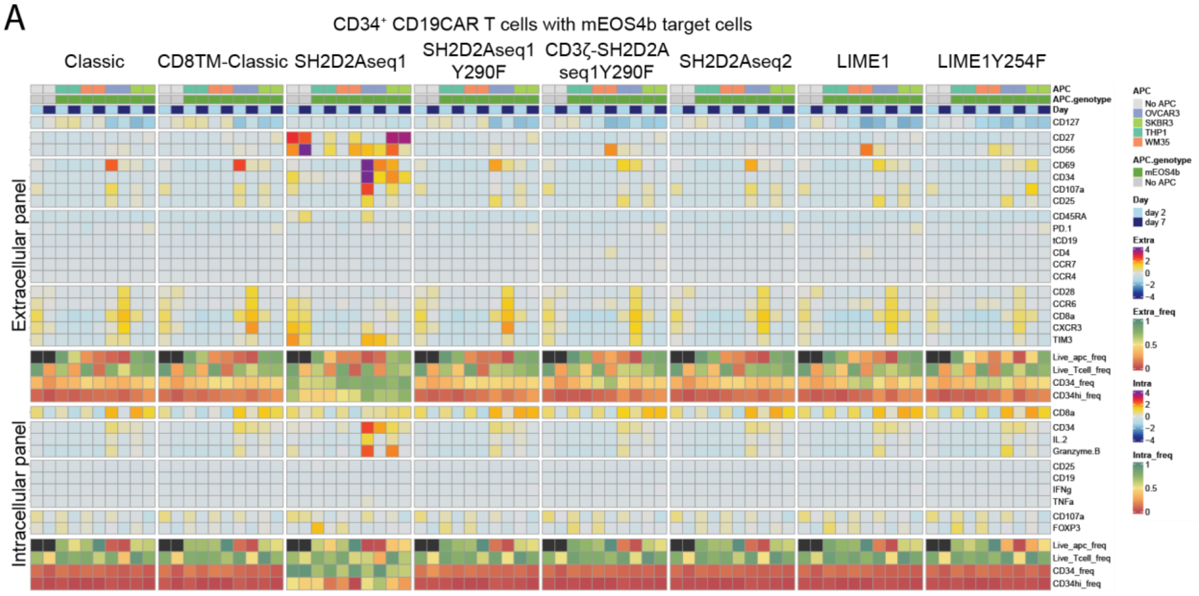

#### Supplementary figure 6

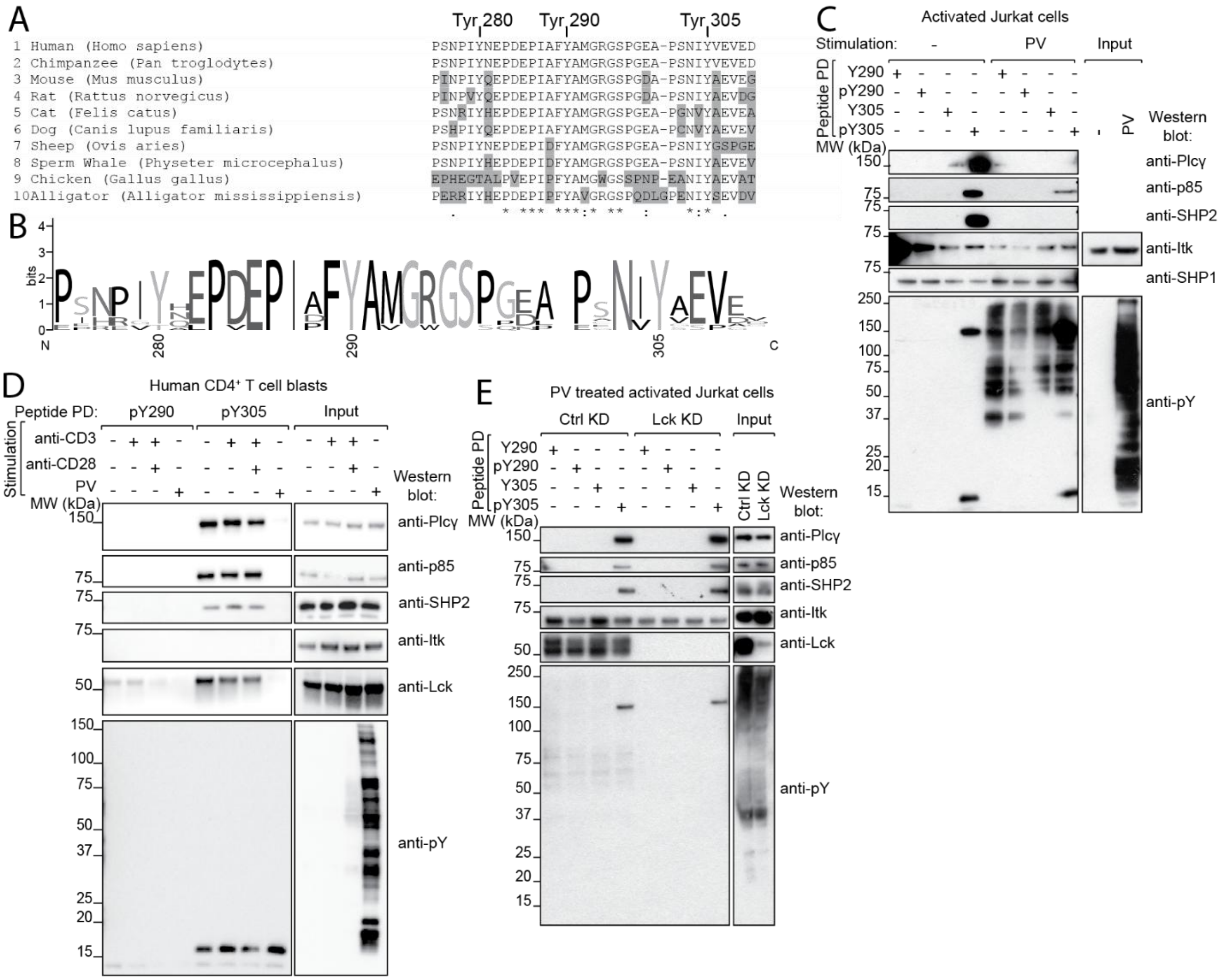

### Supplementary figures legends

#### Supplementary figure 1

**Lck-binding motifs from Lck adaptors molecules are intrinsically disordered**

IDR (intrinsically disordered region) prediction analysis of the Lck adaptor proteins included in this study was performed using AIUPred (https://iupred.elte.hu/) and UniProt identifiers. In each plot, the red line indicates the predicted disorder tendency of each residue, while the blue line shows the context-dependent probability that a residue is part of a binding region. Below the plots, annotated tracks indicate: PFAM (regions from the Protein Families database), EXP DIS (experimentally validated disordered regions), ELM (motifs from the Eukaryotic Linear Motif database), and PTM (post-translational modifications; phosphorylation sites obtained from PhosphoSitePlus). Black brackets denote the protein fragments used in this study.

#### Supplementary figure 2

**Lck adaptors sequences affect surface expression of CAR constructs**

A) Gating strategy used in flow cytometry throughout this project to identify successfully transduced $CAR^{+}$ T cell population.
B) Histograms of surface expression of CD19-CARs with Lck adaptors sequences expressed in JE6.1 T cells from one representative experiment.
C) Histograms of surface expression of CD19-CARs with Lck adaptors sequences expressed in human $CD8^{+}$ T cells from one representative experiment.
D) Summary graph of surface expression of CD34 marker in $CD34^{+}$ JE6.1 T cells transduced with CD19-CARs with Lck adaptors sequences. Each dot represents separate experiment (n=3-8). All data was normalized to classic CD19-CAR measured within given experiment.
E) Summary graph of surface expression of CD34 marker in $CD34^{+}$ human $CD8^{+}$ T cells transduced with CD19-CARs with Lck adaptors sequences. Each dot represents separate experiment (n=2-9) performed in n=2-5 donors. All data was normalized to classic CD19-CAR measured within given experiment.
F) JE6.1 cells were stably transduced with CD19-CARs with Lck adaptors sequences and sorted. Cells were treated for 5 min with pervanadate, lysed with LDS 0,1%/Triton 1% buffer, and incubated for 1h with protein-L covered magnetic beads. PD and lysate samples were analyzed with western blotting.
G) Histograms of surface expression of CD8TM CD19-CARs with Lck adaptors sequences expressed in JE6.1 T cells from one representative experiment.

H) Histograms of surface expression of CD8TM CD19-CARs with Lck adaptors sequences expressed in human $CD8^+$ T cells from one representative experiment.
I) Summary graph of surface expression of CD34 marker in $CD34^+$ JE6.1 T cells transduced with CD8TM CD19-CARs with Lck adaptors sequences. Each dot represents separate experiment (n=2-4). All data was normalized to classic CD19-CAR measured within given experiment.
J) Summary graph of surface expression of CD34 marker in $CD34^+$ human $CD8^+$ T cells transduced with CD8TM CD19-CARs with Lck adaptors sequences. Each dot represents separate experiment (n=4-12) performed in n=1-5 donors. All data was normalized to classic CD19-CAR measured within given experiment.

#### Supplementary figure 3

**Functional phenotype performance of human T cells with CARs with Lck adaptor sequences.**

A) Heatmap presents the comparison between CD19-CAR $(classic)^-$ T cells population within experiment presented in Fig. 2A. These T cells were also present in the samples incubated with the four different target cell lines expressing CD19 or not.
B) CAR T cells with Lck adaptor sequences vs target cell co-culture assay. Heatmap presents the comparison between various CD19-CAR (with Lck adaptor sequences) T cells incubated with the four different target cell lines which do not express CD19.

#### Supplementary figure 4

**Certain phosphotyrosines, when present in CAR, can affect its expression**

A) Histograms of surface expression of CD19-CARs with Lck adaptors sequences after CD3ζ domain expressed in JE6.1 T cells from two representative experiments.
B) Histograms of surface expression of CD19-CARs with Lck adaptors sequences after CD3ζ domain expressed in human $CD8^+$ T cells from one representative experiment.
C) Summary graph of surface expression of CD34 marker in $CD34^+$ JE6.1 T cells transduced with CD8TM CD19-CARs with Lck adaptors sequences after CD3ζ domain. Each dot represents separate experiment (n=2-5). All data was normalized to classic CD19-CAR measured within given experiment.
D) Summary graph of surface expression of CD34 marker in $CD34^+$ human $CD8^+$ T cells transduced with CD8TM CD19-CARs with Lck adaptors sequences after CD3ζ domain. Each dot represents separate experiment (n=2-12) performed in n=2-5 donors. All data was normalized to classic CD19-CAR measured within given experiment.

E) Histograms of surface expression of SH2D2Aseq1 CD19-CARs with mutated phosphotyrosines expressed in JE6.1 T cells from one representative experiment.
F) Histograms of surface expression of CD3ζ-SH2D2Aseq1 CD19-CARs with mutated phosphotyrosines expressed in JE6.1 T cells from one representative experiment.
G) Histograms of surface expression of SH2D2Aseq1 CD19-CARs with mutated phosphotyrosines expressed in human $CD8^+$ T cells from one representative experiment.
H) Histograms of surface expression of CD3ζ-SH2D2Aseq1 CD19-CARs with mutated phosphotyrosines expressed in human $CD8^+$ T cells from one representative experiment.
I) Summary graph of surface expression of CD34 marker in $CD34^+$ JE6.1 T cells transduced with SH2D2Aseq1 CD19-CARs with mutated phosphotyrosines. Each dot represents separate experiment (n=2-5). All data was normalized to classic CD19-CAR measured within given experiment.
J) Summary graph of surface expression of CD34 marker in $CD34^+$ human $CD8^+$ T cells transduced with SH2D2Aseq1 CD19-CARs with mutated phosphotyrosines. Each dot represents separate experiment (n=2-6) performed in n=2-4 donors. All data was normalized to classic CD19-CAR measured within given experiment.
K) JE6.1 cells were stably transduced with CD19-CARs with Lck adaptors sequences and sorted. Cells were treated for 5 min with pervanadate, lysed with LDS 0,1%/Triton 1% buffer, and incubated for 1h with protein-L covered magnetic beads. PD and lysate samples were analyzed with western blotting.
L) Histograms of surface expression of LIME1 and CD3ε CD19-CARs with mutated phosphotyrosines expressed in JE6.1 T cells from one representative experiment.
M) Histograms of surface expression of LIME1 and CD3ε CD19-CARs with mutated phosphotyrosines expressed in human $CD8^+$ T cells from one representative experiment.
N) Summary graph of surface expression of CD34 marker in $CD34^+$ JE6.1 T cells transduced with LIME1 and CD3ε CD19-CARs with mutated phosphotyrosines. Each dot represents separate experiment (n=3-5). All data was normalized to classic CD19-CAR measured within given experiment.
O) Summary graph of surface expression of CD34 marker in $CD34^+$ human $CD8^+$ T cells transduced with LIME1 and CD3ε CD19-CARs with mutated phosphotyrosines. Each dot represents separate experiment (n=2-7) performed in n=2-5 donors. All data was normalized to classic CD19-CAR measured within given experiment.

#### Supplementary figure 5

**Mutation of SH2D2A Tyr290 in CD19-CAR with SH2D2Aseq1 abolishes its rewired signaling.**

A) Altered CAR T cells with SH2D2A or LIME1 sequences vs target cell co-culture assay. Heatmap presents the comparison between various altered CD19-CAR (with SH2D2A or LIME1) T cells incubated with the four different target cell lines which do not express CD19.

#### Supplementary figure 6

**SH2D2A Tyr290 is not recognized by phosphotyrosine binding proteins.**

A) Alignment of TSAd C-terminal tyrosines motifs across ten species. Grey color indicates deviation from human sequence.
B) Protein motif conservation logo generated from sequences in A).
C) JTag T cells, pre-activated with PMA/IO, were treated with PV (5min) or left untreated before being lysed and subjected to PD with indicated TSAd peptides. PD samples were immunoblotted with the indicated antibodies. One representative immunoblot is shown (n=2).
D) Similar to C, but with human $CD4^{+}$ T cell blasts that prior to lysis were treated with anti-CD3 antibody (2min), anti-CD3/anti-CD28 antibodies (2min) or PV (5min). One representative immunoblot is shown (n=2).
E) Similar to C, but JTAg T cells were treated with Lck or control siRNA prior to procedure. One representative immunoblot is shown (n=3).
